# TFIIB-related factor 1 is a nucleolar protein that promotes RNA polymerase I-directed transcription and tumour cell growth

**DOI:** 10.1101/2022.02.20.481212

**Authors:** Juan Wang, Qiyue Chen, Xin Wang, Shasha Zhao, Huan Deng, Baoqiang Guo, Cheng Zhang, Xiaoye Song, Wensheng Deng, Shihua Zhang, Tongcun Zhang, Hongwei Ni

**Affiliations:** College of Life Science and Health, Wuhan University of Science and Technology, Wuhan, 430065, China; School of Materials and Metallurgy, Wuhan University of Science and Technology, Wuhan, 430081, China; Faculty of Biology, Medicine, and Health, University of Manchester, Manchester, M13 9PL, UK; School of Healthcare Science, Manchester Metropolitan University, Manchester, M1 5GD, UK

**Keywords:** BRF1, nucleoli, Pol I transcription, rDNA promoter, tumour cell proliferation.

## Abstract

RNA polymerase I (Pol I) products play pivotal roles in ribosomal assembly, protein synthesis and cell growth. Dysregulation of Pol I product synthesis closely correlates with tumourigenesis. However, the factors and pathways governing Pol I-dependent transcription remain to be identified. Here, we report that TFIIB-related factor 1 (BRF1), a subunit of TFIIIB required for Pol III-mediated transcription, is a nucleolar protein and modulates the rRNA synthesis directed by Pol I. We show that BRF1 can be co-localized with nucleolar protein markers in several human cell types. BRF1 expression positively correlates with Pol I product levels and tumour cell growth *in vitro* and *in vivo*. Mechanistically, BRF1 binds to the Pol I transcription machinery factors and is recruited to the rDNA promoter with them. Alteration of BRF1 expression affected the recruitment of Pol I transcription machinery at the rDNA promoter, the rDNA promoter activity and expression of TBP and TAF1A, indicating that BRF1 modulates Pol I-direct transcription by controlling the expression of selective factor 1 subunits. Collectively, we demonstrate that BRF1 acts as a positive factor to regulate Pol I-directed transcription, suggesting that BRF1 can concurrently modulate Pol I and Pol III transcription and acts as a key coordinator between them.

## Introduction

Ribosomal RNAs (rRNAs) synthesis by RNA polymerase I accounts for over 60% of total transcription activity in eukaryotic cells. Together with the 5S rRNA transcribed by RNA polymerase III (Pol III), cellular rRNA molecules act as essential components of a ribosome to regulate ribosome biosynthesis and translation (*Grummt, 2003; Russell and Zomerdijk, 2005; Goodfellow and Zomerdijk, 2013*). Dysregulation of Pol I transcription causes a range of genetic diseases and cancers (*Drygin et al., 2010; Hannan et al., 2013; Sharifi and Bierhoff, 2018; Ferreira et al., 2020*). Transcriptional initiation by RNA Pol I requires upstream-binding factor (UBF), selective factor 1 (SL1), transcription initiation factor 1A (TIF-1A) and Pol I to assemble at the rDNA promoter, where they form the pre-initiation complex (*Grummt, 2003; Russell and Zomerdijk, 2005*). Apart from general transcription factors, Pol I-dependent transcription is also tightly controlled by oncogenic factors, tumour repressors, signalling factors, chromatin remodelling factors and long noncoding RNAs (*Zhai and Comai 2000; Arabi et al., 2005; Stefanovsky et al., 2006; Zhou et al., 2009; Xing et al., 2009*). It has been reported that nuclear mitotic apparatus protein NuMA modulates rDNA transcription by mediating a nucleolar stress response in a p53-independent manner (*Jayaraman et al., 2017*). Oncogenic protein LYAR (ly1 antibody reactive clone) may enhance rDNA transcription by recruiting bromodomain-containing protein 2 and histone acetyltransferase KAT7 at the rDNA loci (*Izumikawa, et al., 2019*). It has been shown that LncRNA PAPAS (promoter and pre-rRNA antisense) directly interacts with the rDNA enhancer and represses rRNA synthesis through recruiting CHD4/NuRD at the rDNA promoter (*Zhao et al., 2018*). A cellular SUMOylation system can down-regulate the expression of c-MYC and BRF1, and subsequently inhibiting rDNA transcription (*Peng et al., 2019*). Recently, a Che-1*/*AATF (Che-1) protein complex has been shown to bind to the RNA polymerase machinery and maintains rDNA gene transcription (*Sorino et al., 2020*). Cockayne syndrome group A and B proteins (CSA and CSB) regulate transcription of ribosomal DNA (rDNA) genes and ribosome biogenesis by inducing ubiquitination of nucleolin (*Okur et al., 2020*). Despite advance in Pol I-directed transcription, the pathways and factors controlling Pol I-directed transcription remain to be identified.

Pol I general transcription factor IIIB is consisted of TBP, BRF1/2 and BDP1 subunits (*Colbert and Hahn, 1992*); among these subunits, BRF1 is required for the transcription of *5S rRNA* and *tRNA* genes, while BRF2 is essential for transcription of the *U6 RNA* gene (*Roberts et al., 1996; Yeganeh and Hernande, 2020*). Deregulation of TFIIIB expression is closely associated with cancer development (*White, 2004*; *Yeganeh and Hernande, 2020*). During transcriptional initiation by Pol III, TFIIIC binds to the internal box A and B within tRNA genes, subsequently recruiting TFIIIB and Pol III at the transcription start site (TSS) (*Grewal, 2015*; *Yeganeh and Hernande, 2020*). It has been shown that tumour repressors, including p53, RB and Maf1, can associate with TFIIIB to prevent the assembly of the Pol III transcription machinery and repress Pol III-dependent transcription (*Chesnokov et al., 1996; Sutcliffe et al., 2000; Goodfellow et al., 2008; White, 2008*). Conversely, oncogenic protein c-MYC targets TFIIIB to activate Pol III-directed transcription (*Gomez-Roman et al., 2003; White, 2008*). Additionally, signal pathways such as RAS/ERK, PI3K/AKT and JNK indirectly regulate Pol III gene transcription (*Moir and Willis 2013; Felton-Edkins et al., 2003, Sriskanthadevan-Pirahas et al., 2018; Zhong et al., 2009; Zhong et al., 2013; Woiwode et al., 2008*). Aberrantly high expression of TFIIIB subunits has been observed in human cancer cells (*Daly et al., 2005; Zhong et al., 2004*). For example, breast cancers and hepatocellular carcinomas have abnormally high expression of BRF1, suggesting that BRF1 is an excellent biomarker for cancer diagnosis (*Huang et al., 2019; Fang et al., 2017; Lei et al., 2017*). Previous and recent evidences show that BRF1 acts as a positive factor to regulate the Pol III transcription (*Zhong et al., 2009; Peng et al., 2020)*.

When we previously investigated the role of FLNA in Pol III-directed transcription (*Peng et al., 2020; Wang at al*., 2016), BRF1 was accidently observed to be present in the nucleoli of human osteosarcoma cells (see *Figure 1-figure supplement*). However, whether BRF1 is a nucleolar protein in other cell types and what role BRF1 plays in the nucleoli remained unknown. In this study, we show that BRF1 is quite abundant in the nucleoli of several tumour cell types, and BRF1 positively regulates in Pol I-directed transcription and tumour cell growth *in vitro* and *in vivo*. We further dissected the mechanism by which BRF1 modulates Pol I-mediated transcription using combined techniques.

## Results

### BRF1 is abundantly present in the nucleoli of human cell lines

As described above, BRF1 is a subunit of TFIIIB required for Pol III-directed gene transcription (*Colbert and Hahn, 1992*, *Roberts et al., 1996*). Unexpectedly, the co-localization between BRF1 and filamin A (FLNA) was observed in the nucleoli of SaOS2 cells (*Figure 1-figure supplement A*) when we investigated the role of FLNA in Pol III-mediated transcription. Our recent work revealed that both BRF1 and fibrillarin (FBL, a nucleolar protein marker) were also co-localized in the nucleoli of SaOS2 cells (*Figure 1-figure supplement B*). Based on these data, we supposed that BRF1 could also be a nucleolar protein although it was found in the nucleoplasm and plays an important role in Pol III transcription (*Peng et al., 2020; Wang at al*., 2016). To substantiate this supposition, we first performed immunofluorescence (IF) assays using HeLa cells and the antibodies against BRF1, FLNA and nucleophosmin (NPM1), respectively. Intriguingly, the co-localization between BRF1 and FLNA (or between BRF1 and NPM1) was observed in HeLa cell nucleoli, indicating that BRF1 is present in the nucleoli of HeLa cells (*Figure 1A and B*). We next determined whether this observation could be reproduced when nucleolar particles purified from HeLa cells were used for IF assays. As illustrated in *Figure 1C and D*, either NPM1 or FBL could be co-localized with BRF1 in nucleolar particles. We subsequently examined the presence of BRF1 in the nucleolar particles and other cellular fractions by Western blot. Immunoblotting data showed that BRF1 was able to be detected in the nucleolar fraction and had a stronger signal than that in the other cellular fractions, indicating that BRF1 abundantly exists in the nucleoli of HeLa cells (*Figure 1E*). Next, we addressed whether BRF1 also exists in normal human cells. To this end, we performed IF assays using human primary cervical epithelial cells (ATCC, PCS-480-011, it is here abbreviated into HuCEC). *Figure 1F* shows that BRF1 could be observed in the nucleoli of HuCEC cells. Taken together, we confirm that BRF1 is not only present in the nucleoli of tumour cells but also in those of normal human cells. These results suggest that BRF1 could play a role in human cell nucleoli.

**Figure 1.**
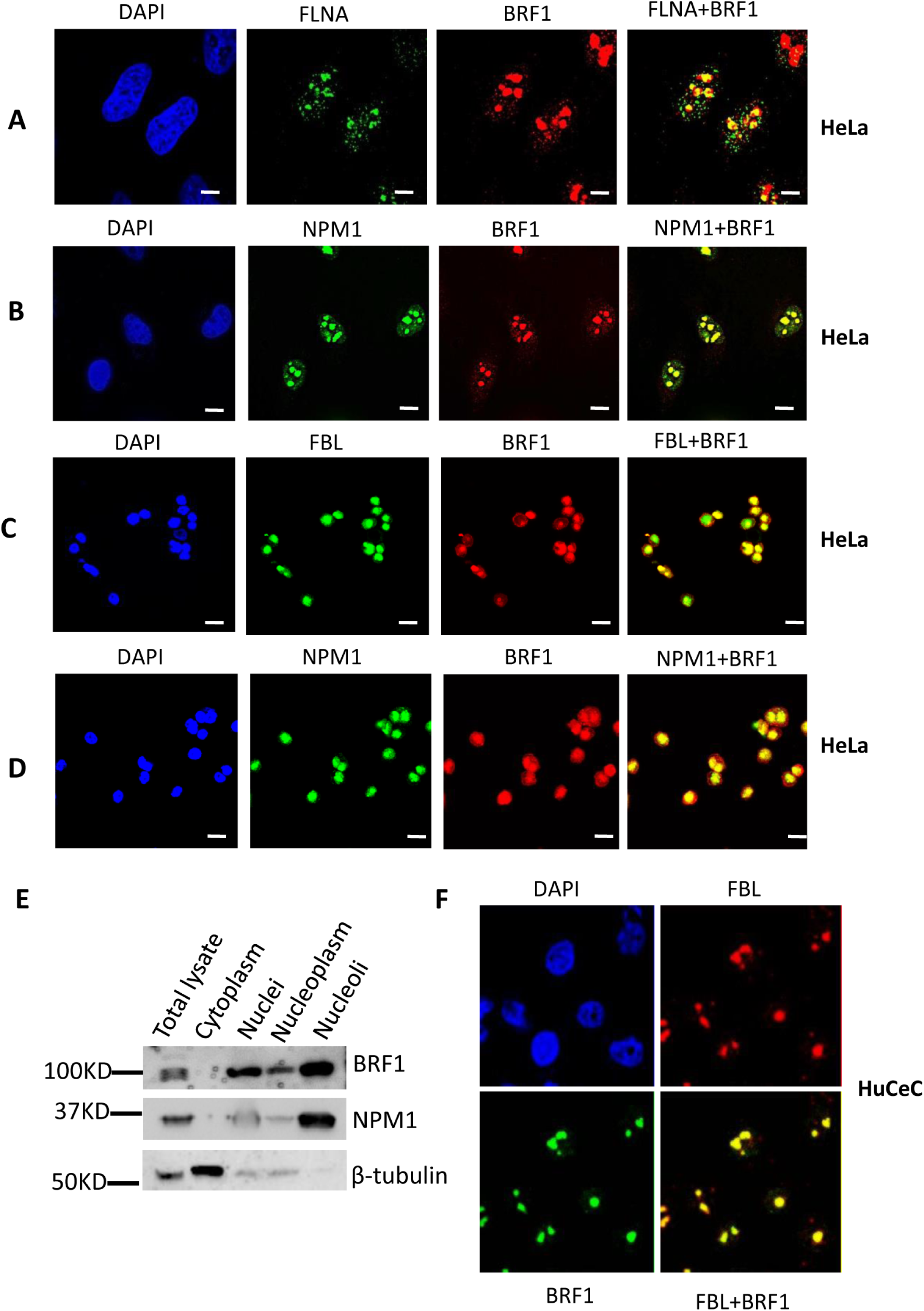
BRF1 is localized to the nucleoli in HeLa cells. (A) BRF1 and FLNA were co-localized in the nucleoli of HeLa cells. Immunofluorescence (IF) staining was performed using HeLa cells and the antibodies against BRF1 or FLNA. The specimens from IF assays were observed under a DeconVision fluorescent microscope and imaged with a 60x objective lens (Olympus). The scale bars in the images represent 2 μm. (**B**) BRF1 and NPM1 were co-localized in the nucleoli of HeLa cells. IF staining was performed as described in *A* in which the NPM1 antibody replaced the FLNA antibody. The scale bars in the images represent 5 μm. (**C**) Analysis of the co-localization between BRF1 and FBL in the nucleoli particles purified from HeLa cells. Nucleoli were purified from the nuclei of HeLa cells through sonication and gradient centrifugation. IF assays were performed using nucleolar samples and the antibodies against BRF1 or FBL. The IF samples were observed under a confocal fluorescent microscope and imaged with a 100x object lens (Olympus). The scale bars in the images represent 1 μm. (**D**) Analysis of the co-localization between BRF1 and NPM1 in the nucleoli particles purified from HeLa cells. IF staining was performed as for **C** in which the FBL antibody was replaced by the NPM1 antibody. The scale bars in the images represent 1 μm. (**E**) Immunoblotting analysis for the nucleoli fraction and other cellular fractions obtained from HeLa cells. Nucleoli fraction was obtained as described in **C**. Western blot was performed using the antibodies as indicated in the panel. (**F**) The co-localization analysis between BRF1 and FBL in human cervical epithelial cells (HuCEC). IF assay was performed with human cervical epithelial cells and the antibodies as indicated. The specimens were observed and imaged under a confocal fluorescence microscope. The scale bars in the images represent 2 μm. **Figure 1-figure supplement**: BRF1 is present in the nucleoli within SaOS2 cells. **Figure 1-source data**

### BRF1 acts as a positive factor to regulate Pol I-directed transcription in transformed cell lines

The eukaryotic nucleolus is an important subcellular structure that is responsible for 45S pre-rRNA synthesis, rRNA processing and initial assembly of ribosomes. Considering that BRF1 was found in the nucleolus of human cells, we next determined whether it is required for the synthesis of 45S pre-rRNA. We first transfected BRF1 siRNA or control siRNA into HeLa cells, and immunofluorescence assays were performed using the cells transfected and BRF1 antibody. Data showed that BRF1 siRNA transfection reduced BRF1 localization to the nucleoli of HeLa cells (*Figure 1-figure supplement C*), further confirming the presence of BRF1 in HeLa cell nucleoli. Next, expression of BRF1 and pre-rRNA in these cells was monitored by RT-qPCR or Western blot. Noticeably, BRF1 siRNA transfection reduced BRF1 expression (*Figure 2A and B*) and the synthesis of Pol I products. As expected, 5S rRNA expression, a positive control (a Pol III product), was also inhibited by BRF1 siRNA transfection (*Figure 2C*). Meanwhile, a comparable result was obtained in 293T cells transfected with BRF1 siRNA or control siRNA (*Figure 2-figure supplement 1A-C*). However, we failed to get results from SaOS2 cells transfected with BRF1 siRNA due to the low efficiency of transient transfection. These data indicate that BRF1 down-regulation could repress in Pol I-mediated transcription. We then determined whether the result obtained above could be reproduced in the cell lines stably expressing BRF1 shRNA or control shRNA generated by a lentiviral infection system. Both RT-qPCR and Western blot data showed that the cell lines stably expressing BRF1 shRNA or control shRNA, including HeLa, SaOS2, 293T and M2, were obtained (*Figure 2D, E, G and H*; *Figure 2-figure supplement 1D, E, G and H*). Analysis of Pol I products by RT-qPCR revealed that BRF1 silencing in these cell lines dampened the expression of Pre-rRNA, 18S rRNA and 28S rRNA (*Figure 2F and I*; *Figure 2-figure supplement 1F and I*), indicating that BRF1 down-regulation, indeed, inhibits expression of Pol I products. We next examined whether BRF1 expression change could affect the synthesis of nascent rRNA using EU (5-ethynyle uridine) assays, where the rRNA incorporated by EU was analysed using an EU detection kit (Beyotime, China). Strikingly, BRF1 silencing in HeLa cells inhibited EU incorporation in the nascent rRNA (*Figure 2J and K*), suggesting that BRF1 may activate ribosomal RNA gene expression.

**Figure 2.**
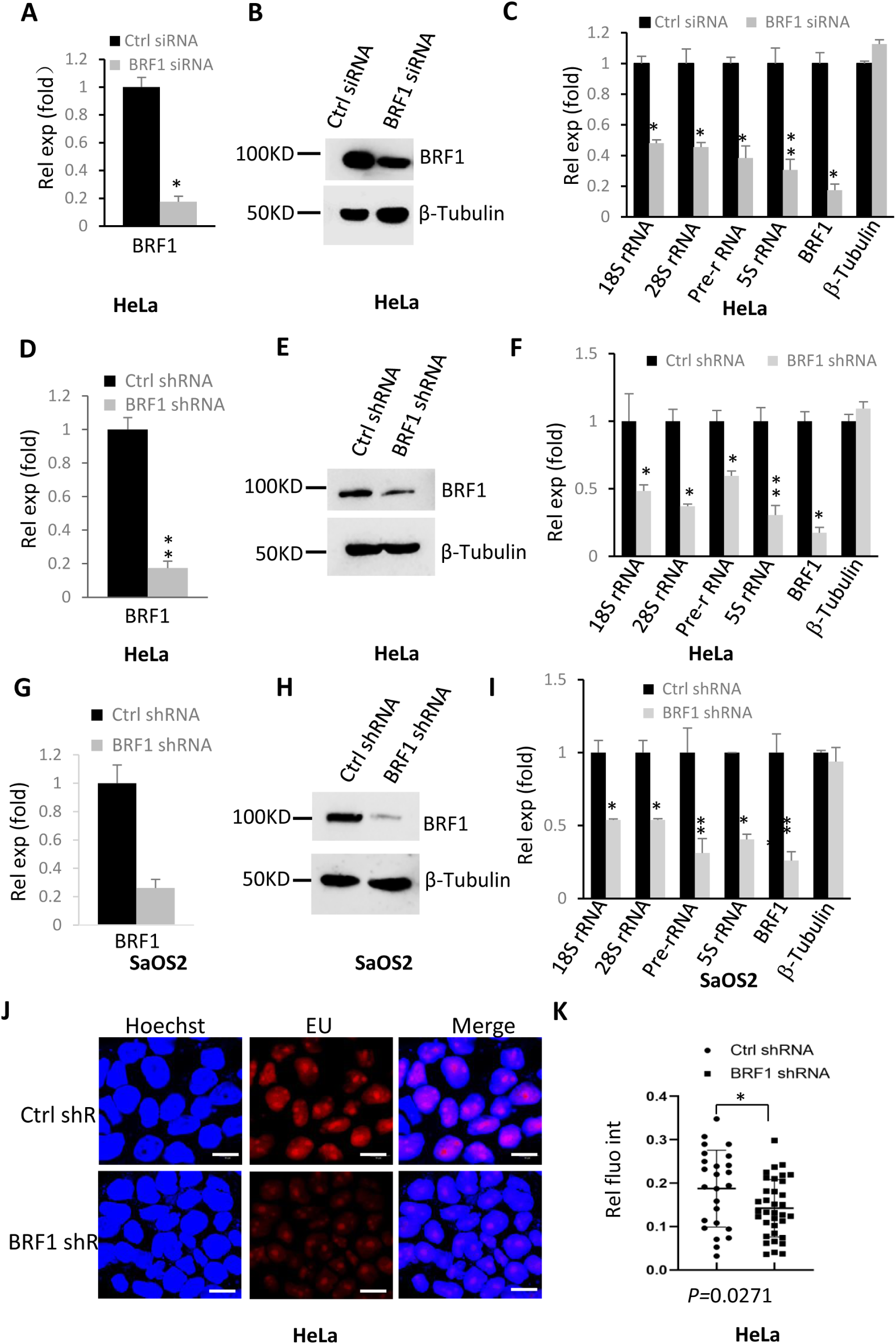
BRF1 knockdown inhibited Pol I-directed gene transcription. (**A**-**C**) Transfection of BRF1 siRNA reduced Pol I-directed transcription in HeLa cells. HeLa cells were transfected with BRF siRNA; BRF1 mRNA and protein expression were monitored by RT-qPCR (**A**) and Western blot, respectively (**B**). Ribosomal RNA gene expression was analysed by RT-qPCR (**C)**. (**D**, **E**) Generation of HeLa cell lines stably expressing BRF1 shRNA or control shRNA. HeLa cells were infected with lentiviral particles expressing BRF1 shRNA or control shRNA. The cell lines stably expressing BRF1 shRNA or control shRNA were screened by 96-well plates and determined by RT-qPCR (**D**) and Western blot (**E**). (**F**) BRF1 shRNA stable expression inhibited Pol I-mediated transcription in HeLa cells. Total RNA was extracted from the cell lines obtained in **D** and **E** and rRNA gene expression was analysed by RT-qPCR. (**G**-**I**) BRF1 depletion by shRNA repressed Pol I transcription in SaOS2 cells. SaOS2 cell lines stably expressing BRF1 shRNA or control shRNA were generated using a lentiviral infection system and determined by RT-qPCR (**G**) and Western blot (**H**). Pol I products were monitored by RT-qPCR (**I**). (**J** and **K**) A EU assay result showing the effect of BRF1 depletion on rRNA synthesis. HeLa cell lines stably expressing BRF1 shRNA or control shRNA were labelled with 5-ethynyl uridine (EU). The specimens from EU assays were observed and imaged with a fluorescence microscope (**J**) and the red fluorescence was analysed with the ImageJ software (**K**). The scale bar in each image represents 50 μm. Each column in **A**, **C**, **D**, **F**, **G** and **I** represents the mean±SD of three biological replicates (n=3). *, *P*≦0.05; **, *P*≦0.01, *P* values were obtained by one way ANOVA. **Figure 2-figure supplement 1.** BRF1 knockdown reduced Pol I-mediated transcription. **Figure 2-figure supplement 2.** Effect of BRF1 overexpression on Pol I–directed transcription. **Figure 2-figure supplement 3.** BRF1 overexpression enhanced the rRNA levels newly synthesized in the nulceoli of HeLa cells. **Figure 2-figure supplement 4.** BRF1 promotes Pol I-mediated transcription and proliferation of HuCEC cells **Figure 2-source data; Figure 2-figure supplement 1-source data; Figure 2-figure supplement 2-source data, Bar or point graphs-source data**

To verify the positive role of BRF1 in Pol I–mediated transcription, we transfected HeLa and 293T cells using the vectors expressing HA-BRF1 and its control vectors (pcDNA). Expression of BRF1 and 45S rRNA was monitored by Western blot and RT-qPCR, respectively. We show that BRF1 overexpression enhanced rRNA gene expression, including Pol III product 5S rRNA expression (*Figure 2-figure supplement 2A-D*). We then generated HeLa, 293T and M2 cell lines stably expressing HA-BRF1 and their control cell lines using a lentiviral infection system. Western blot data illustrated that the cell lines stably expressing HA-BRF1 were successfully established (*Figure 2-figure supplement 2E, G and I*). As expected, stable overexpression of BRF1 could activate Pol I-directed transcription in these cell types (*Figure 2-figure supplement 2F, H and J*). These results are consistent with those obtained in transient transfection assays. We then examined the effect of BRF1 overexpression on the rRNA newly synthesized by performing EU assays. We show that BRF1 overexpression in HeLa enhanced EU-labelled rRNA levels (*Figure 2-figure supplement 3A and B*), indicating that BRF1 overexpression stimulated the synthesis of Pol I products. Interestingly, the result from the analysis of Pol I products using the normal cell lines (HuCEC) with BRF1 depletion or overexpression is consistent with that obtained from tumour cells (*Figure 2-figure supplement 4A and B*). Taken together, these results indicate that BRF1 can act as an activator to modulate Pol I-directed transcription in human cell lines.

### BRF1 can independently and directly regulate Pol I-directed transcription

We have examined the effect of BRF1 expression alteration on Pol I-directed transcription using indirect methods, including RT-qPCR and EU labelling. Next, we verified the result obtained above using Northern dot blot. The total RNA was extracted from HeLa cell nuclei, and Northern dot blot assays were performed using the resulting RNA and the probes detecting pre-rRNA. Notably, BRF1 silencing reduced pre-rRNA synthesis (*Figure 3A and B*); whereas BRF1 overexpression enhanced pre-rRNA products (*Figure 3C and D*). These data further confirmed that BRF1 could positively modulate Pol I-directed transcription. BRF1 was originally found to be one of TFIIIB subunits, which positively regulates Pol III-directed transcription (*Peng et al., 2020*). We next examined whether BRF1 regulates Pol I product expression independence of Pol III product change. HeLa cells with BRF1 silencing or overexpression and their control cells were cultured in the presence or absence of 50 μM ML-60218 (a Pol III transcription inhibitor), and Pol I-directed transcription was detected by RT-qPCR. Interestingly, the presence of ML-60218 neither significantly affected the activation of Pol I-directed transcription caused by BRF1 overexpression nor the inhibition of Pol I-directed transcription caused by BRF1 silencing. Although a little impact from Pol III inhibitor (ML-60218) was observed, the effect is not significant (*Figure 3E and F*). In contrast, expression of 5S rRNA, one of Pol III products, was severely affected by the presence of ML-60218. These data suggest that BRF1 can independently regulate Pol I-directed transcription rather than depending on the change of Pol III products. To determine whether BRF1 directly regulates Pol I-directed transcription, we performed *in vitro* transcription assays using the rDNA promoter-driving reporter vectors and the HeLa nuclear extract with or without BRF1 depletion by BRF1 antibody. Data showed that BRF1 depletion reduced the transcription activity of the rDNA promoter (*Figure 3G and H*). Considering that transcription *in vitro* were performed in a cell-free system, where a small amount of Pol III products in the reaction system supposed to be stable and couldn’t interfere with the assays, the result from *in vitro* transcription assays indicates that BRF1 can independently and directly regulate Pol I-directed transcription.

**Figure 3.**
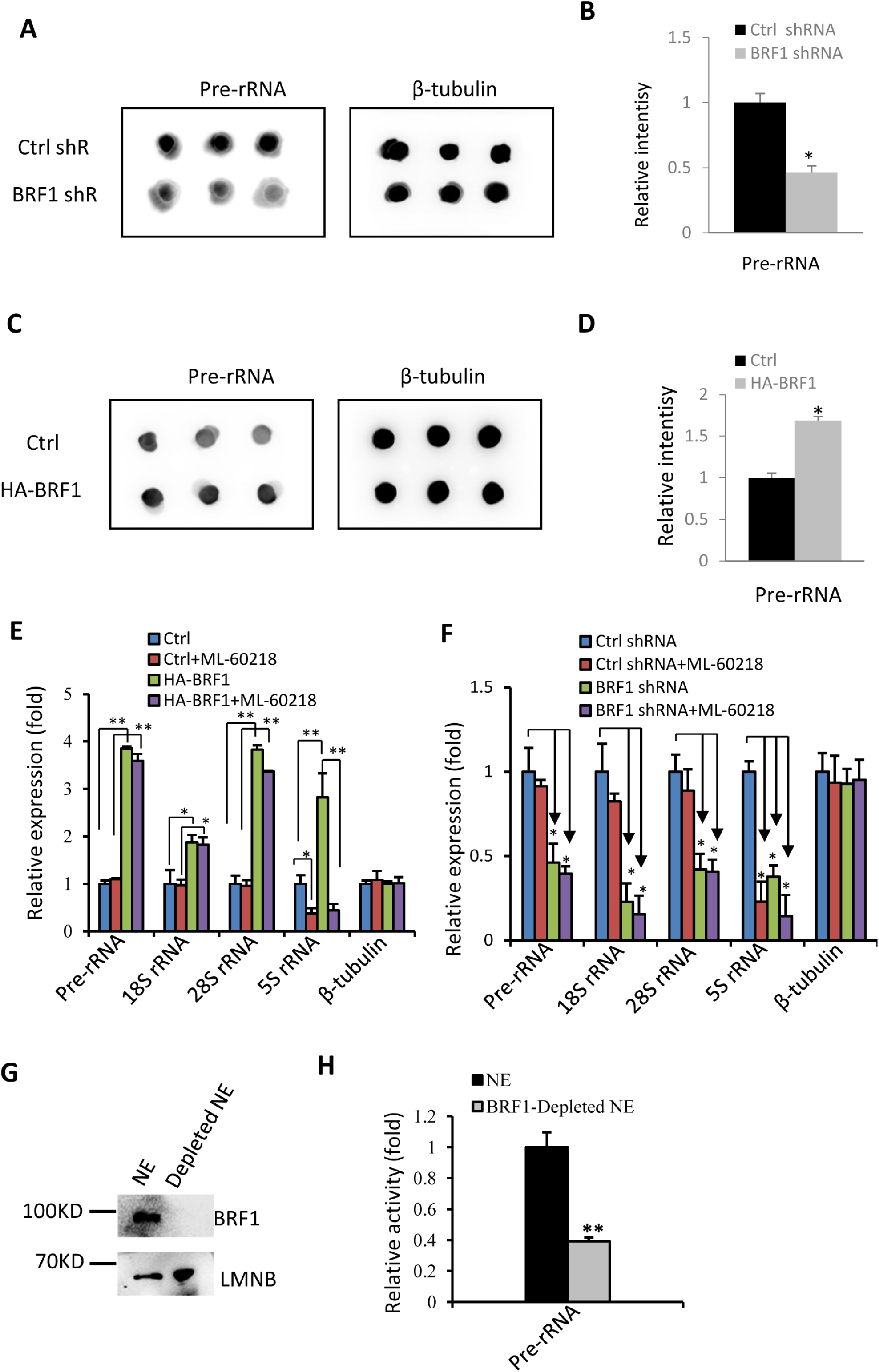
BRF1 can directly regulate Pol I-directed transcription. (**A**) A Northern dot blot result for 45S pre-RNA (left panel) and β-tubulin mRNA from HeLa cells expressing BRF1 shRNA and control shRNA. (**B**) The quantified result of Northern dot blots obtained in **A**. (**C**) A Northern dot blot result for 45S pre-RNA (left panel) and β-tubulin mRNA from a HeLa cell line expressing HA-BRF1 shRNA and its control cell line. (**D**) The quantified result of Northern dot blots obtained in (**C**). (**E**) Pol III-specific inhibitor did not significantly affect the activation of Pol I transcription caused by BRF1 overexpression. RT-qPCR was performed using the RNA extracted from HeLa cell lines as indicated in the absence or presence of 50 μM ML-60218. (**F**) Pol III-specific inhibitor did not significantly aggravate the inhibition of Pol I transcription caused by BRF1 silencing. RT-qPCR was performed using the RNA extracted from HeLa cell lines as indicated in the absence or presence of 50 μM ML-60218. (**G**) A Western blot result showing the HeLa nuclear extract with or without BRF1 depletion. (**H**) *In vitro* transcription assays for the rDNA promoter-driving reporter gene, where the reporter gene (*luciferase*) transcribed *in vitro* was detected by RT-qPCR. Each column in **B**, **D**-**F** and **H** represents the mean±SD of three biological replicates (n=3). *, *P*≦0.05; **, *P* ≦0.01, *P* values were obtained by one way ANOVA. **Figure 3-source data, Bar or point graphs-source data**

### Pol I product change caused by BRF1 up- and down-regulation influenced cell proliferation in vitro and in vivo

Pol I product levels closely correlate with cell growth and proliferation. Thus, it is necessary to determine if alteration of Pol I products mediated by BRF1 contributed cell proliferation. The effect of BRF1 depletion on cell proliferation was first examined using cell counting and MTT (3-[4,5-dimethylthiazol-2-yl]-2,5 diphenyl tetrazolium bromide) assays. We show that BRF1 silencing inhibited transformed cell proliferation, including HeLa, M2, SaOS2 and 293T (*Figure 4A and B*; *Figure 4-figure supplement 1*). To confirm this result, we performed EdU (5-ethynyl-2′-deoxyuridine) assays using HeLa cell lines, where the rate of EdU-labelled positive cells was calculated. We show that BRF1 knockdown reduced the rate of the EdU-labelled cells when compared to the control cell line, confirming that BRF1 promotes HeLa cell proliferation (*Figure 4C and D*). Using the same strategy, we examined the effect of BRF1 overexpression on cell proliferation. The data from cell counting and MTT assays showed that BRF1 overexpression stimulated cell proliferation activity, including HeLa, 293T and M2 cells (*Figure 4E and F*; *Figure 4-figure supplement 2*). A consistent result was obtained by EdU assays when HeLa cells with BRF1 overexpression were examined (*Figure 4G and H*). Additionally, we show that BRF1 expression positively correlated with proliferation activity in normal human cells such as HuCEC cells (*Figure 2-figure supplement 4C and D*). To determine whether the activation of Pol I-directed transcription caused by BRF1 overexpression contributed the increase of cell proliferation, we performed cell proliferation assays by adding a Pol I-specific inhibitor CX-5461 into HeLa cells expressing HA-BRF1. Data showed that the presence of CX-5461 inhibited cell proliferation and Pol I-mediated transcription in HA-BRF1-expressing cells when compared to control cells or the HA-BRF1-expressing cells in the absence of CX-5461 (*Figure 4I*; *Figure 4-figure supplement 3*). These results indicate that the activation of Pol I-mediated transcription caused by BRF1 overexpression contributed the increase of cell proliferation although the contribution of Pol III product increase cannot be excluded.

**Figure 4.**
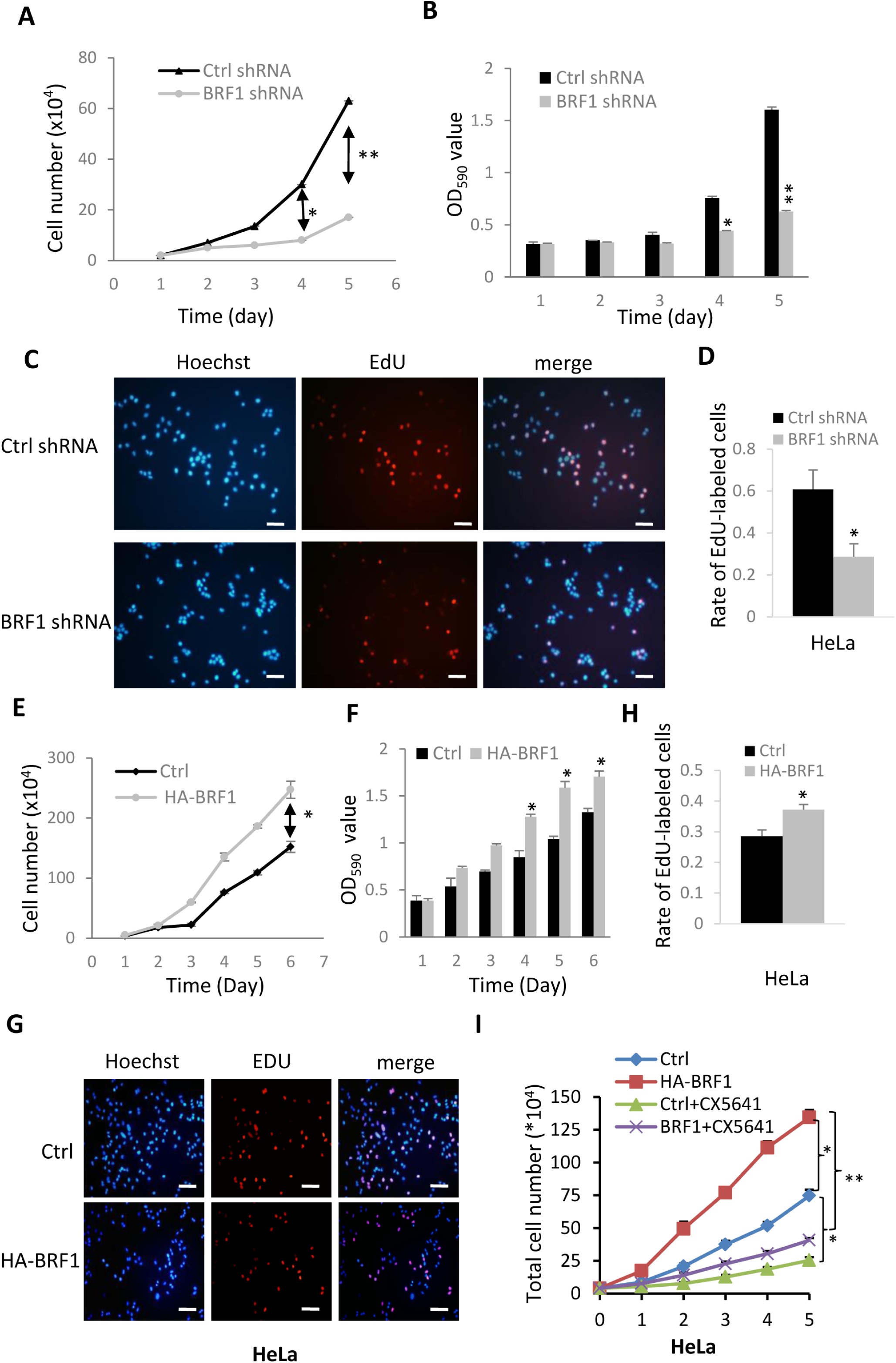
BRF1 promotes cell proliferation by increasing Pol I transcription. (**A**, **B**) Knockdown of BRF1 repressed HeLa cell proliferation. HeLa cell lines stably expressing BRF1 shRNA or control shRNA were seeded in 12-well and 96-well plates. Cell proliferation was analysed using the cell counting method (**A**) and the MTT assays (**B**). (**C**, **D**) Analysis of HeLa cell proliferation by the EdU assays. HeLa cell lines stably expressing BRF1 shRNA or control shRNA seeded in 12-well plates were labelled with EdU and the effect of EdU labelling was monitored by using the EdU detection kit. The specimens were observed and imaged with a fluorescence microscope (**C**). The EdU–labelled cells were counted under a microscope and analysed with the GraphPad 6.0 (**D**). The scale bar in the image represents 200 μm. (**E**, **F**) BRF1 overexpression enhanced HeLa cell proliferation. A HeLa cell line stably expressing HA-BRF1 and its control cell line were seeded in 12-well and 96-well plates. Cell proliferation was analysed using the cell counting method (**E**) and the MTT assays (**F**). (**G**, **H**) EdU assays for the HeLa cell line with BRF1 overexpression and its control cell line.A HeLa cell line stably expressing HA-BRF1 and its control cell line were grown in 12-well plates. EdU assays were performed as done for (**C**) and (**D**). The EdU-labelled cells were imaged (**G**) and statistically analysed (**H**). The scale bar in the image represents 200 μm. (**I**) CX-5461 suppressed the increased activity of cell proliferation induced by BRF1 overexpression. A HeLa cell line expressing HA-BRF1 and its control cell line were grown in 12 well plates, 4 μg of CX-5461 were added into each well containing the HA-BRF1-expressing cells. The cells were harvested at each time point and used for cell counting or Po I product analysis. Each column in **A**, **B**, **D**-**F**, **H** and **I** represents the mean±SD of three biological replicates (n=3). *, *P*≦0.05; **, *P*≦0.01, *P* values were obtained by one way ANOVA. **Figure 4-figure supplement 1.** BRF1 knockdown inhibited proliferation activity for transformed cell lines. **Figure 4-figure supplement 2.** BRF1 overexpression stimulated proliferation activity for transformed cell lines. **Figure 4-figure supplement 3.** CX-5461 inhibited the activation of Pol I transcription caused by BRF1 overexpression in HeLa cells. **Figure 2-figure supplement 4.** BRF1 promotes Pol I-mediated transcription and proliferation of HuCEC cells. **Bar or point graphs-source data**

We have shown that BRF1 up- and down-regulation promoted tumour cell proliferation *in vitro*. Whether this result can be observed *in vivo* remained unclear. To address this question, we performed tumour formation assays by injecting a HeLa cell line expressing BRF1 shRNA or its control cell line into nude mice (n=10 for each group). *Figure 5A* shows that the sizes of the tumours during tumour growth were reduced by BRF1 knockdown. Analysis of tumour weight showed that the weights of tumours were also significantly decreased by BRF1 silencing (*Figure 5B and C*), indicating that BRF1 down-regulation inhibited tumour growth *in vivo*. Hematoxylin and eosin staining showed that the tumour tissue had large and condensed nuclei, which were distinct from the cervical tissue or the skin tissues (*Figure 5D*). Immunohistochemistry and immunoblotting analysis revealed that tumour tissues retained the original features of the tumour cell lines before injection (*Figure 5E and F*). *Figure 5G* shows that the tumour tissues expressing BRF1 shRNA had lower Pol I transcription activity than the control shRNA-expressing tissues. These data indicate that the inhibition of Pol I-mediated transcription caused by BRF1 down-regulation can lead to the repression of tumour cell growth *in vivo* but the contribution of Pol III products cannot be excluded.

**Figure 5.**
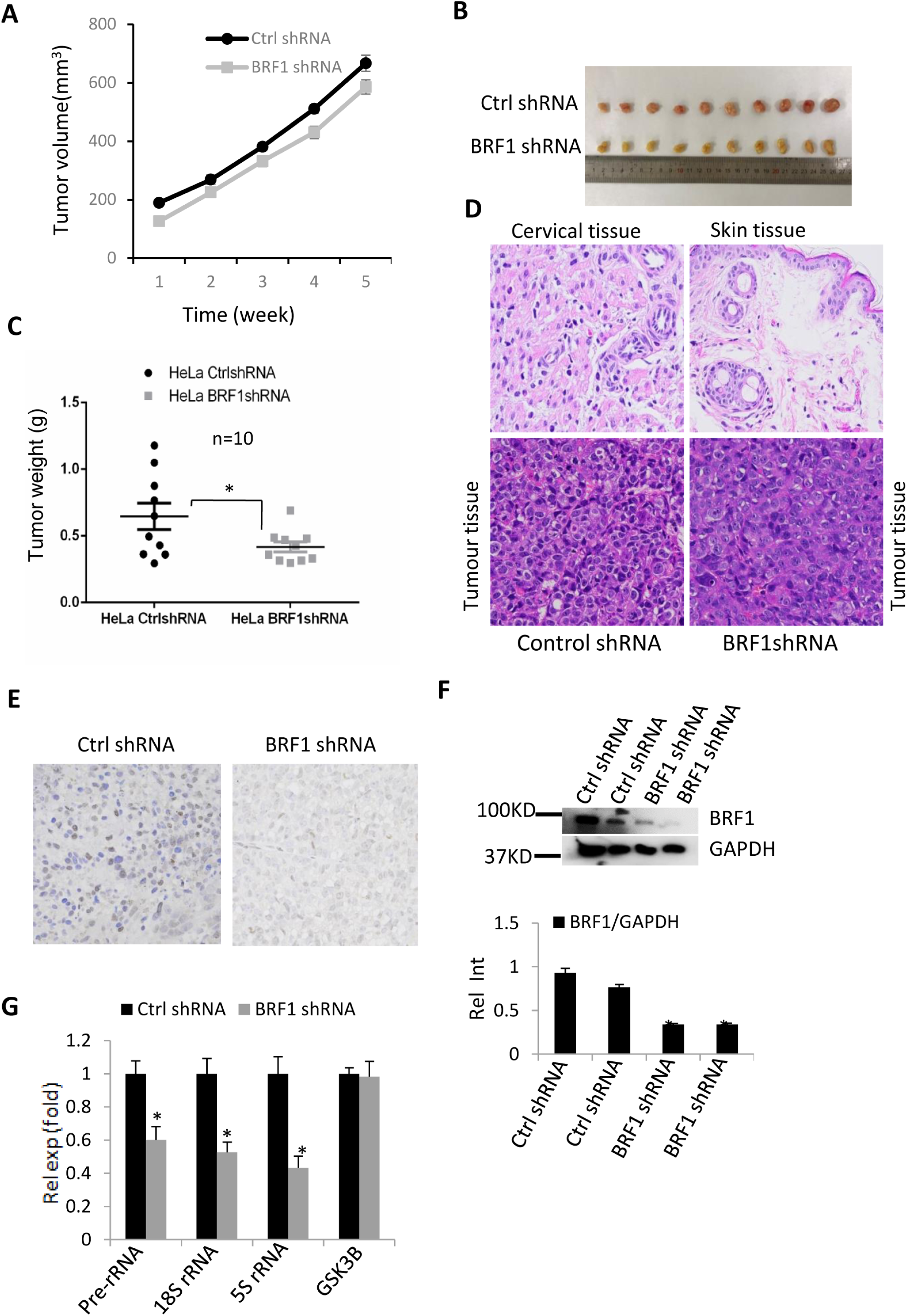
BRF1 knockdown repressed tumour growth *in vivo*. (**A**) Comparison of the tumour volumes between the HeLa cells expressing BRF1 shRNA and the HeLa cells expressing control shRNA during tumour formation. HeLa cells stably expressing BRF1 shRNA or control shRNA were subcutaneously injected into nude mice (n=10 for each group). At one week after injection, the sizes of tumours were measured every 3 days using a Vernier caliper. The volume of individual tumours was subjected to statistical analysis. (**B**) An image showing the difference of the tumour sizes between the tumour samples expressing BRF1 shRNA and the tumour samples expressing control shRNA. Nude mice were euthanized after injection for 5 weeks. The tumour samples from the mice were carefully taken out of the mice and photographed using a digital camera (Canon). (**C**) BRF1 knockdown inhibited tumour growth. Tumour samples obtained in B were weighed on an electronic scale and the data were analysed with the GraphPad 6. (**D**) Representative images showing the HE staining results of the tumour, cervix, and skin tissues from nude mice. The specimens were observed and imaged using an upright light microscope (Olympus). The scale bars in the images represent 100 μm. (**E**) Immunohistochemistry staining of the tumour tissues expressing BRF1 shRNA or control shRNA using BRF1 antibody. The scale bars in the images represent 100 μm. (**F**) Immunoblotting analysis of the tissue samples expressing BRF1 shRNA or control shRNA. The intensity of the Western blot bands (the upper panel) was analysed by the ImageJ software (the bottom panel). (**G**) Analysis of Pol I products for the tumour tissue expressing BRF1 shRNA or control shRNA. Each column in **G** represents the mean±SD of three biological replicates (n=3) *, *P*≦0.05; **, *P*≦0.01, *P* values were obtained by one way ANOVA. **Figure 5-source data, Bar or point graphs-source data**

### BRF1 binds to the Pol I transcription machinery

To understand how BRF1 modulates Pol I-directed transcription, we examined the connection between BRF1 and the Pol I transcription machinery using combined techniques. Immunofluorescence (IF) assays were firstly performed using HeLa cells and the antibodies against BRF1 or the factors from the Pol I transcription machinery. IF staining showed that BRF1 was able to be co-localized with TIF-1A, RPA40 and UBF in the nucleoli of HeLa cells (*Figure 6A and B*; *Figure 6-figure supplement 1A*). Next, IF staining was performed using nucleolar particles purified from HeLa cells. Data showed that the co-localization between BRF1 and the factors within the Pol I transcription machinery such as TIF-1A, RPA43 and TBP could be observed in the nucleolar particles (*Figure 6C and D*; *Figure 6-figure supplement 1B*), suggesting that the interaction between BRF1 and the Pol I transcription machinery factors is possible. To determine if BRF1 interacts with these factors, we performed co-immunoprecipitation (co-IP) assays using BRF1 antibody. *Figure 6E* shows that the BRF1 antibody was able to precipitate TIF-1A, UBF, TAF1A, TBP and RPA43 proteins from the HeLa nuclear extract. In reciprocal co-IP assays, BRF1 protein could be precipitated by the antibodies against the factors as indicated (*Figure 6F and G*). To verify the authenticity of BRF1 co-IP result, we tested if BRF1 can bind to SL1 other subunits and Pol III transcription factors by performing co-IP assays using the antibodies against BRF1, TAF1B and GTF3C2, respcetively. Western blot showed that BRF1 could also associate with TAF1B and GTF3C2 proteins (*Figure 6-figure supplement 2*). Taken together, co-IP results indicate that BRF1 can bind to components of the Pol I transcription machinery.

**Figure 6.**
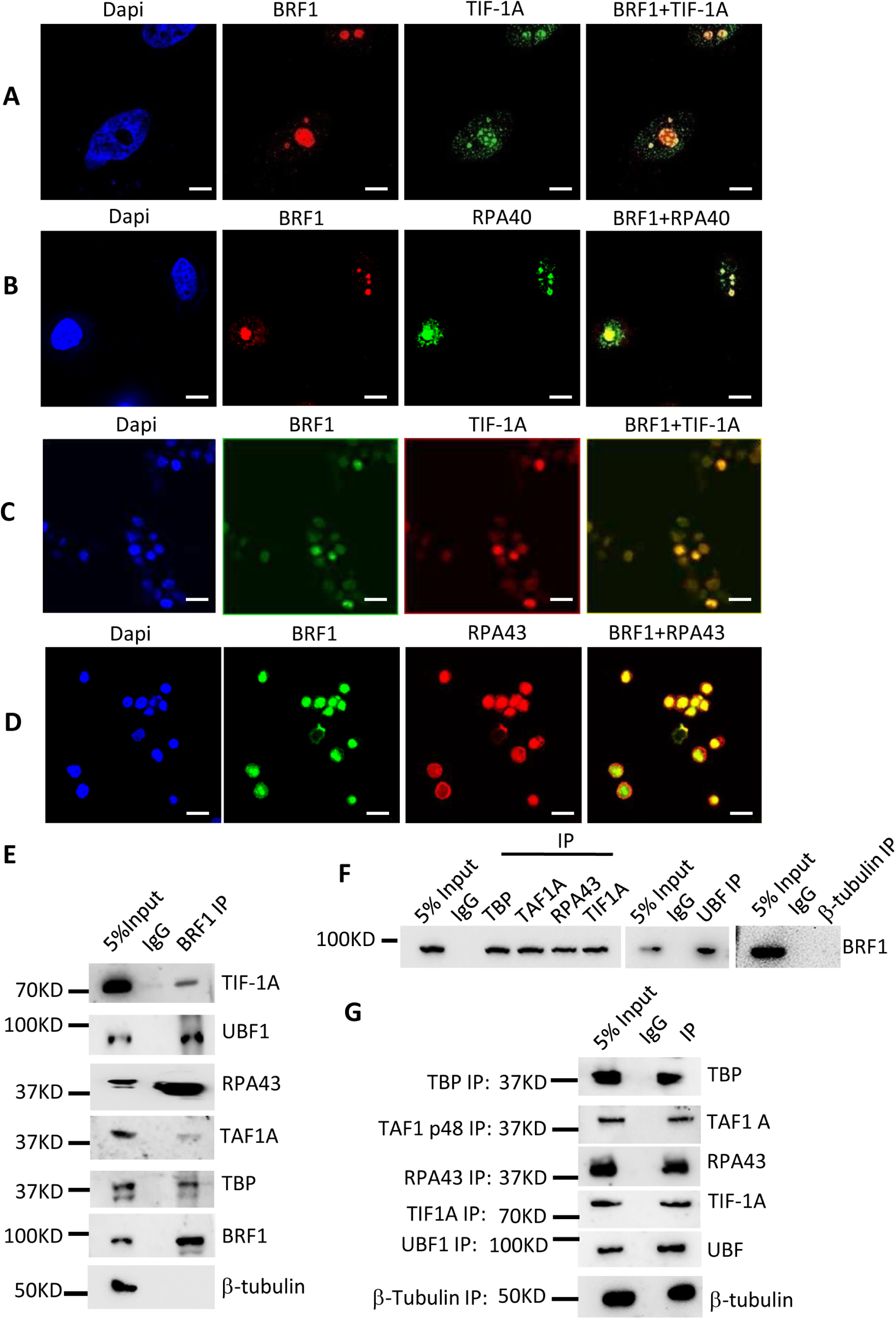
BRF1 interacts with components of Pol I transcription machinery. (A) BRF1 and TIF-1A were co-localized in the nucleoli of HeLa cells. IF staining was performed using the BRF1 and TIF-1A antibodies. The scale bar in each image represents 2.5 μm. (**B**) BRF1 and RPA40 were co-localized in the nucleoli of HeLa cells. IF staining was performed using BRF1 antibody and RPA40 antibody. The scale bar in each image represents 2.5 μm. (**C**) The co-localization analysis between BRF1 and TIF-1A using the nucleolar particles purified from HeLa cells. The scale bar in each image represents 2.5 μm. (**D**) The co-localization analysis between BRF1 and RPA43 using the nucleoli particles purified from HeLa cells. The scale bar in each image represents 5 μm. (**E**) Components of the Pol I transcription machinery could be precipitated by the BRF1 antibody. HeLa nuclear extract was prepared and used for BRF1 IP assays. BRF1-binding proteins in the IP samples were detected by Western blot using the antibodies as indicated in the panel. (**F**, **G**) BRF1 could be precipitated by the antibodies against Pol I-related proteins. IP assays were performed with the antibodies against TBP, TIF-1A, TAF1A, UBF, and RPA43, respectively. BRF1 (**F**) and proteins as indicated (**G**) were detected by Western blot. **Figure 6-figure supplement 1.** The co-localization analysis between BRF1 and Pol III transcription factors in HeLa cells. **Figure 6-figure supplement 2.** BRF1 binds to TFIIIC subunit GTF3C2 and SL1 subunit TAF1B. **Figure 6-source data, Figure 6-figure supplement 2-source data**

### BRF1 binds to the rDNA promoter and modulates the assembly of the Pol I transcription machinery at the promoter

To gain the details about how BRF1 regulates Pol I-directed transcription, we determined whether BRF1 bound to the rDNA promoter by performing ChIP assays. *Figure 7A* shows that BRF1 had a significant occupancy at the rDNA promoter in HeLa cells, although its occupancy at the rRNA-coding region and the intergenic spacer was significantly reduced. Since BRF1 binds to the Pol I transcription machinery, we next determined if BRF1 could bind to the rDNA promoter along with components of the Pol I transcription machinery by performing sequential ChIP assays. As demonstrated in *Figure 7B*, BRF1 occupied the rDNA promoter in company with UBF, TIF-1A, TBP and RPA43. Considering that BRF1 binds to the rDNA promoter, we next determined if alteration of BRF1 expression affected the rDNA promoter activity. The rDNA promoter was cloned into the pGL3-basic reporter vector (*Figure 7C*). The promoter–driving reporter gene vectors were transfected into the HeLa cell lines expressing BRF1 shRNA or HA-BRF1 and their control cell lines. RT-qPCR showed that BRF1 knockdown repressed the rDNA promoter activity. In contrast, BRF1 overexpression activated the rDNA promoter activity, indicating that BRF1 can directly regulate the rDNA promoter activity (*Figure 7D and E*). Interestingly, in the experiments using the vector containing a 45S rRNA cDNA fragment driven by the rDNA promoter (*Figure 7F)*, the result obtained was consistent with that from the reporter assays (*Figure 7G*). To further understand how alteration of BRF1 expression modulates Pol I-directed transcription or the rDNA promoter activity, we performed ChIP assays using HeLa cell lines with BRF1 depletion or overexpression and their control cell lines. The relative occupancies of the Pol I transcription machinery factors at the rDNA promoter were analysed by qPCR. *Figure 7H* illustrates that BRF1 down-regulation reduced the recruitments of TBP, TIF-1A, UBF, TAF1A and RPA43 at the rDNA promoter. In contrast, BRF1 overexpression enhanced the recruitments of these factors at the promoter (*Figure 7I*). These data suggest that BRF1 modulates rDNA transcription by controlling the Pol I transcription machinery assembly at the rDNA promoter.

**Figure 7.**
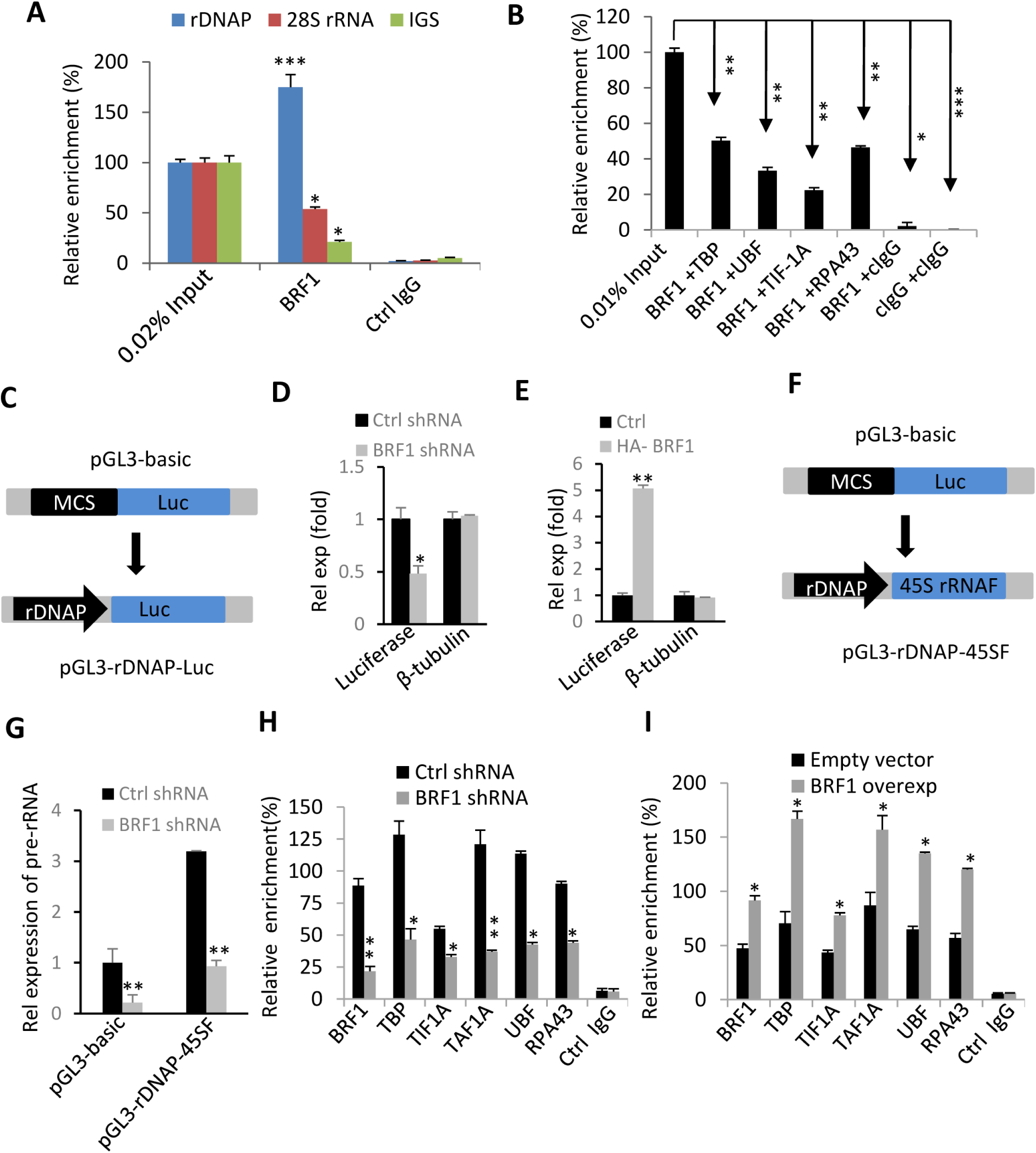
BRF1 modulates Pol I-mediated transcription by controlling the Pol I transcription machinery assembly at the rDNA promoter. (A) A ChIP-qPCR result showing BRF1 binding to the rDNA promoter (rDNAP), coding region (28S rRNA), and intergenic spacer (IGS). ChIP assays were performed using HeLa cells and anti-BRF1 antibody or control IgG. The relative enrichment was obtained by comparing the relative quantity of promoter DNA in 1 μL ChIP sample to that in 0.02% of input DNA. (**B**) Sequential ChIP assays for BRF1 and the factors of the Pol I transcription machinery. One μL of the DNA samples (40 μL) recovered from the sequential ChIP assay or 0.5 ng genomic DNA (0.01% input) was analysed by qPCR. The relative enrichment was obtained as described in **A**. (**C**) A scheme showing the construction of the report vector pGL3-rDNAP-Luc. rDNAP: rDNA promoter, MCS: multiple cloning site. Luc: *luciferase* gene. (**D**) BRF1 knockdown inhibited the rDNA promoter activity. HeLa cell lines as indicated were transfected with the rDNA promoter-driving reporter vectors in **C**. Reporter gene expression was analysed by RT-qPCR using luciferase primers. (**F**) BRF1 overexpression enhanced the rDNA promoter activity. A HeLa cell line expressing HA-BRF1 and its control cell line were transfected with the rDNA promoter-driving reporter vectors in **C**. Reporter gene expression was detected as for **D**. (**F**) A scheme showing the construction of the vector pGL3-rDNA-45SF. 45SF: 45S rRNA fragment. (**G**) BRF knockdown inhibited expression of rRNA fragment driven by the rDNAP. HeLa cell lines as indicated were transfected with the vector pGL3-rDNAP-45SF. Expression of 45S rRNA was performed by RT-qPCR using rRNA gene primers. (**H**) BRF1 knockdown inhibited the recruitments of the Pol I transcription machinery factors at the rDNA promoter. ChIP assays were performed using HeLa cell lines as indicated in the graph. Quantitative PCR was performed using 1 μL of the DNA sample (40 μL) recovered from ChIP assays or 1 ng genomic DNA (0.03% of input DNA). The relative enrichment was obtained as described in **A**. (**I**) BRF1 overexpression enhanced the recuitment of the Pol I transcription machinery factors at the rDNA promoter. ChIP assays were performed using HeLa cell lines and the antibodies as indicated. ChIP assays, quantitative PCR, and data analysis were performed as described in **A**. Each bar in the graphs represents the mean±SD of three biological replicates (n=3)*, *P*≦0.05; **, *P*≦0.01, *P* values were obtained by one way ANOVA. **Bar or point graphs-source data**

### BRF1 regulates expression of TBP and TAF1A and the stability of TAF1A proteins

To understand why the alteration of BRF1 expression influenced the Pol I transcription machinery assembly at the rDNA promoter, we analysed expression of the factors in the Pol I transcription machinery by Western blot using the cell lines with BRF1 knockdown or overexpression and their control cell lines. As shown in *Figure 8A*, BRF1 overexpression enhanced TBP and TAF1A expression but did not affect the expression of TIF-1A, UBF and RPA43. Conversely, BRF1 knockdown inhibited TBP and TAF1A expression but did not affect expression of other factors detected in the assays (*Figure 8B*). These data suggest that BRF1 regulates the Pol I transcription machinery assembly at the rDNA promoter through controlling the expression of selective factor1 (SL1) subunits TBP and TAF1A. Since BRF1 was found to associate with TBP and TAF1A (*Figure 6E-G*), we tested whether BRF1 depletion could affect the stability of TBP and TAF1A proteins. To this end, we performed co-IP assays using TBP and TAF1A antibodies and the HeLa cells transfected with BRF1 siRNA or control siRNA. The protein levels obtained from the co-IP and the ubiquitination of TBP and TAF1A were analysed by Western blot. We show that BRF1 knockdown reduced the precipitation of TBP and TAF1A. These data are consistent with that observed in *Figure 8B*. Interestingly, BRF1 knockdown increased the ubiquitination levels of TAF1A proteins but did not affect that of TBP when compared to the control samples, indicating that BRF1 depletion can influence that stability of TAF1A proteins (*Figure 8C and D*). Taken together, these data suggest that BRF1 modulates the Pol I transcription machinery assembly at the rDNA promoter by influencing expression and stability of the SL1 subunits.

**Figure 8.**
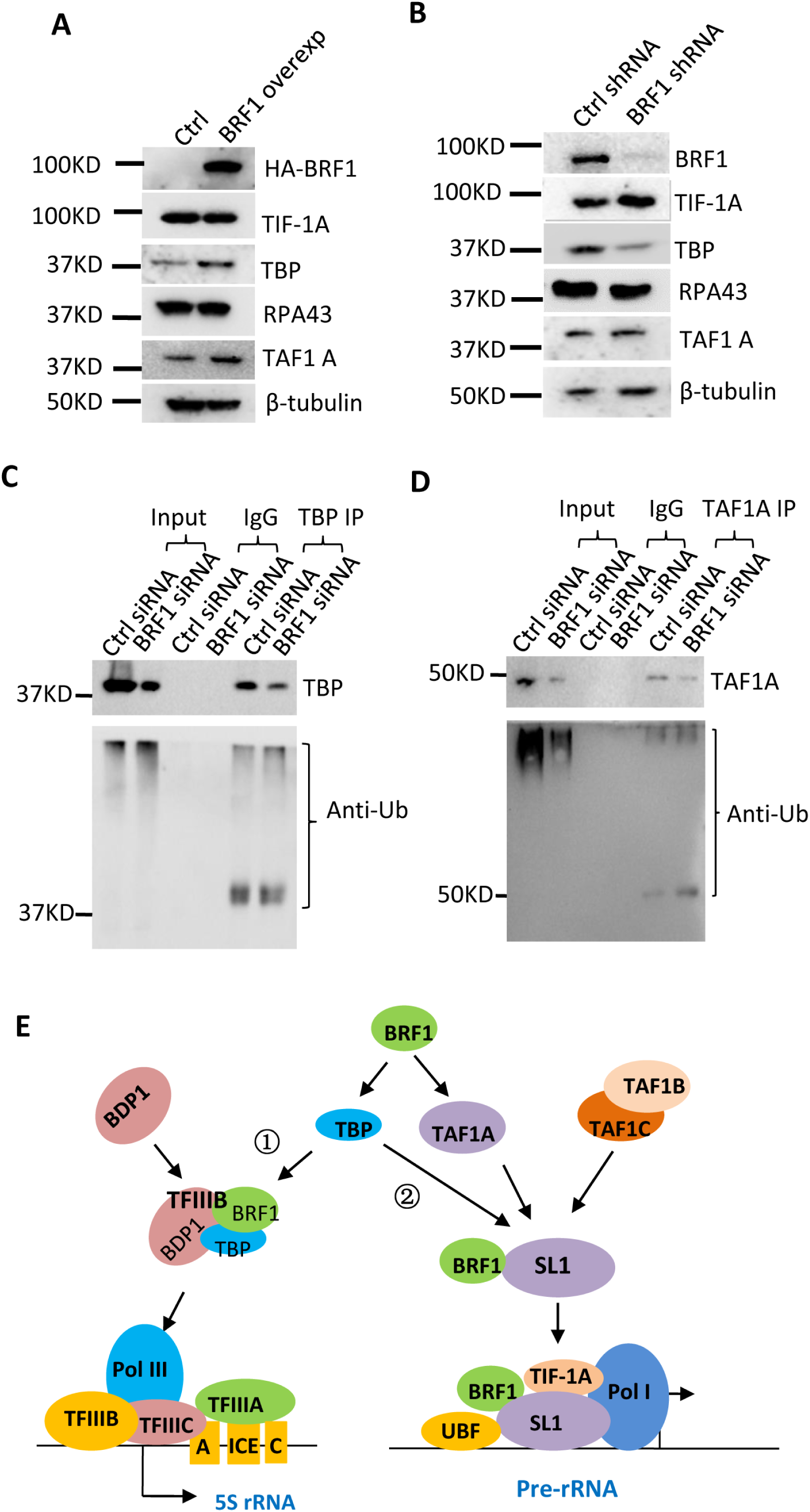
BRF1 regulates the expression of TBP and TAF1A at both protein and mRNA levels. (A) The effect of BRF1 overexpression on the expression of the Pol I transcription machinery factors. A HeLa cell line stably expressing HA-BRF1 and its control cell line were cultured in 6-well plates and harvested for Western blot assays using the antibodies as indicated. (**B**) The effect of BRF1 knockdown on the expression of the Pol I transcription machinery factors. HeLa cell lines stably expressing BRF1 shRNA or control shRNA were cultured and harvested for Western blot. Protein expression was analysed using the antibodies as indicated. (**C**, **D**) BRF1 knockdown affected the ubiquitination of TAF1A proteins but did not impact on that of TBP. IP assays were performed using the antibodies against TBP (**C**) or TAF1A (**D**). Western blot was performed using the antibodies against TBP, TAF1A, and ubiquitin, respectively. *, *P* ≦0.05; **, *P*≦0.01, *P* values were obtained by one way ANOVA. (**E**) A model showing how BRF1 regulates Pol I and III transcriptions. ①, BRF1 binds to TBP and BDP1 to form a TFIIIB so that it regulates Pol III –mediated transcription. ②, The pathway was discovered in this study, where BRF1 promotes the expression of TBP and TAF1A at both protein and mRNA levels, which subsequently enhances the recruitments of SL1 and other factors in the Pol I transcription machinery at the rDNA promoter, eventually controlling Pol I-mediated gene transcription. **Figure 8-source data**

Base on the data obtained in this study, we proposed a model by which BRF1 independently regulates Pol I- and Plo III-directed transcription in human cells. Specifically, BRF1 binds TBP and BDP1 to form the transcription factor IIIB that regulates Pol III transcription (*Figure 8E-*①). Meanwhile, BRF1 associates with TBP, TAF1A, TFF1B and other subunits to form the SL1 complex regulating Pol I transcription (*Figure 8E*-②). Alteration of BRF1 expression affects the protein levels of the SL1 subunits and their occupancies at the rDNA promoter, which accordingly controls the Pol I transcription machinery assembly at the rDNA promoter and Pol I-directed transcription (*Figure 8E*-②).

## Discussion

BRF1 has been identified to be a subunit of TFIIIB and activates Pol III gene transcription (*Colbert and Hahn, 1992; Roberts et al., 1996; Lei et al., 2107; Peng et al., 2020*). In this study, however, we show that BRF1 is abundantly present in the nucleoli of several human cell types (*Figure 1*; *Figure 1-figure supplement*) and can act as a positive factor to modulate Pol I gene transcription (*Figure 2*; *Figure 2-figure supplement 1-3*). Thus, in this study we identified a novel role of BRF1 in Pol I-directed transcription. The findings from the present and previous studies suggest that BRF1 can concurrently modulate both Pol I- and Pol III-mediated transcription by associating with Pol I and Pol III transcription machinery. Although the other factors, including oncogenic factors, tumour repressors, and signalling factors, have been confirmed to regulate both Pol I and Pol III transcriptions (*Sharifi and Bierhoff, 2018; White, 2004; White, 2008*). However, these factors modulate Pol I and Pol III transcription through indirectly regulating the activities of general transcription factors (*White, 2008*). The data from this work suggest that BRF1 acts as a direct link between Pol I and Pol III transcription so that human cells co-ordinates Pol I- and Pol III-dependent transcription by controlling cellular BRF1 expression levels.

We show that knockdown of BRF1 inhibited proliferation activity for several human cell types. Conversely, BRF1 overexpression enhanced cell proliferation activity (*Figure 4*; *Figure 4-figure supplement 1-3*). Additionally, the BRF1 down-regulation reduced the sizes and weights of tumours derived from the HeLa tumour cells xenografted into nude mice (*Figure 5*). These data indicate that BRF1 can promote tumour cell growth *in vitro* and *in vivo*. Previous studies has shown that BRF1 is abnormally high expression in alcohol-associated cancers, including hepatocellular carcinoma and breast cancer (*Huang et al., 2019; Fang et al., 2017; Lei et al., 2017*, *Zhong et al., 2016*), suggesting that BRF1 can be applied to a potential biomarker for cancer diagnosis in the clinic and a therapeutic target of alcohol-associated cancers (*Huang et al., 2019; Fang et al., 2017; Lei et al., 2017*). Indeed, tamoxifen has already been found to repress alcohol-induced BRF1 expression and Pol III transcription activity in ERα^+^ breast cancer (*Zhong et al., 2014*). A recent study has shown that the compound betaine impedes the activation of both BRF1 expression and Pol III transcription induced by alcohol and subsequently inhibits tumour cell growth (*Deng et al., 2014*). In this study, we found that BRF1 promoted cell proliferation by stimulating Pol I transcription activity, whereas a pol I inhibitor CX-5461 severely repressed the increase of cell proliferation and the activation of Pol I-directed transcription induced by BRF1 overexpression. This finding suggests that anti-cancer drugs may be developed by inhibiting BRF1, Pol I and Pol III transcription activities.

We demonstrate that BRF1 associated with the Pol I transcription machinery factors and bound to the rDNA promoter along with these factors (*Figure 6; Figure 7A and B*). Alteration of BRF1 expression influenced the rDNA promoter activity and the occupancies of the Pol I transcription machinery factors at the promoter (*Figure 7C–I*). Additionally, we found that the alteration of BRF1 expression also affected the expression of TBP and TAF1A proteins (*Figure 8A and B*). These data suggest that BRF1 modulates Pol I-directed transcription by affecting the activities of SL1 subunits. Since BRF1 binds to TBP, TAF1A and TAA1B (*Figure 6E-G*; *Figure 6-figure supplement 2*), which are subunits of SL1 required for Pol I transcription (*Russell and Zomerdijk, 2005*). Thus, it is plausible that alteration of BRF1 expression affected the expression of TBP and TAF1A. BRF1 silencing also affected the stability of TAF1A, suggesting that BRF1 modulate Pol I transcription assembly by influencing SL1 stability. Taken together, we identified a novel function for BRF1 in Pol I-directed transcription, suggesting that BRF1 can independently modulate Pol- and Pol III-mediated transcription. The findings from this study provide novel insights into the mechanism of Pol I-directed transcription and the mechanism of the coordination between Pol I- and Pol III-directed transcription.

## Experimental procedures

### Plasmids, cells and reagents

The pLV-U6-EGFP-Puro lentiviral plasmid was purchased from Inovogen Tech Co. (Beijing, China). Three different cDNA fragments encoding BRF1 shRNAs were cloned downstream of the U6 promoter within the plasmid. The pGL3-basic plasmid containing a reporter gene was purchased from Promega Co. SaOS2, HeLa, 293T and M2 cell lines were obtained from ATCC. and cultured in their respective media with the supplement of 10% fetal bovine serum (Biowest Co.) and 1× penicillin/streptomycin (Hyclone Co.). DNA and RNA extraction kits were purchased from Axygene Co. Biological reagents, including restriction enzymes, reverse transcription enzyme, PCR, and transfection reagents, were obtained from Thermo Fisher Scientific Co. All other chemicals were purchased from Sigma-Aldrich (Merck).

### Immunofluorescence assays

SaOS2 and HeLa cells were grown on the small round coverslips (14 mm in diameter) in their respective complete media. At 60% confluence, the culture medium was removed and cells were washed twice using 1×PBS solution, followed by fixing for 10 min using 4% formaldehyde freshly prepared with 1×PBS. Thereafter, immunofluorescence (IF) co-localization between BRF1 and nucleolar proteins or Pol I-related factors from the Pol transcription machinery was performed using the method as described previously (*Deng et al., 2012; Wang et al., 2016*). The cell specimens were observed under a Decon Vision fluorescence microscope and imaged with a Z-stack tool and a 60× objective lens (Olympus). Deconvolution for the original images was then performed with the SoftWORx software (GE Healthcare), and the resulting images were analysed with the ImageJ software (NIH).

For IF assays with nucleolar particles, nuclei were isolated from 2×10^7^ HeLa cells by using buffer A (10 mM Hepes pH7.9, 10 mM KCl, 1.5 mM MgCl2, 0.5 mM DTT) and a Dounce tissue homogenizer with a tight pestle. The nuclei were obtained by centrifuging the cell lysate at 1000 rpm (Bekman GS-6 centrifuge, GH-3.8 rotor), then re-suspended with 1 mL S1 solution (0.25 m sucrose, 10 mM MgCl_2_). The nuclei mixtures were slowly added to 2 mL S2 solution (0.35 m sucrose, 10 mM MgCl_2_) and subjected to centrifugation at 2500 rpm using the same centrifuge. The purified nuclei were re-suspended with the S2 solution and subject to disrupting for 6×10 sec using an sonicator (ScientZ-IID, Ningbo, China) with the smallest tip. The resulting nucleoli were purified via a sucrose gradient centrifugation using a S3 solution (0.8 M sucrose, 10 mM MgCl_2_). After centrifugation for 10 min at 3500 rpm, the supernatant was discarded and the nucleolar pellet was suspended in 500 μL S2 solution; followed by centrifuging for 5 min at 2500 rpm. The pellet obtained was re-suspended with 200 μL S2 solution so that highly purified nucleoli were achieved. Twenty microliters of the nucleoli were fixed with 100 μL of 4% formaldehyde-PBS solution. After fixation, the samples fixed were subjected to IF staining using BRF1 antibody and the antibodies against nucleolar proteins or the factors from the Pol I transcription machinery. The specimens were observed under a confocal fluorescence microscope and imaged with a 100× objective lens (Olympus). The primary antibodies used in IF assays were purchased from Abcam (FLNA, ab76289; Fibrillarin, ab166630; Nucleophosmin, ab183340; UBF, ab244287; RPA40, ab196657; RPA43, ab99305) and Santa Cruz Biotech (BRF1, SC-81405). Fluorescent secondary antibodies were from Thermo Scientific (Cat. No. A-11001, A-11005, A-11034 and R37117).

### Transfection, generation of cell lines and rRNA expression analysis

For transient transfection assays, three BRF1 small interfering RNA molecules (siRNAs) were synthesized by Genewiz Co. (Shuzhou, China). HeLa or 293T cells were cultured in 12-well plates for 24 hours prior to transfection assays. Two μL of Turbofect (Thermo Scientific) were mixed with 60 pmol siRNA mixture (20 pmol for each siRNA) and the resulting suspension were mixed into the cells in each well. At 48 hours posttransfection, cells were harvested and used for the analysis of BRF1 expression and Pol I-mediated transcription. To generate cell lines stably expressing BRF1 shRNA or HA-BRF1, we transfected 293T cells using the lentiviral vectors expressing BRF1 shRNA or HA-BRF1 and the packaging vectors pH1 and pH2. After 48 hours, the medium containing lentiviral particles was collected and filtrated with sterile 0.45-μm filters. The medium filtrated was used to infect HeLa, M2 and SaoOS2 cells that were cultured in 12-well plates. The cell line stably expressing BRF1 shRNA or HA-BRF1 were selected by puromycin and screened with 96-well plates. Positive cell lines were determined by immunoblotting with BRF1 antibody. For rRNA expression analysis in the cell line stably expressing BRF1 shRNA or HA-BRF1, RT-qPCR was performed as described previously (*Deng et al., 2012*). For the analysis of rRNA newly synthesized, HeLa cells with BRF1 depletion or overexpression were cultured in 12-well plates and labelled with 5-Ethynyl Uridine (EU) for 2 hours, followed by fixation with a 3.7% formaldehyde solution freshly prepared with 1×PBS. The RNA labelled with EU was detected using the manual provided by the Cell-Light EU Apollo 555 in Vitro Imaging Kit (RiboBio, Guangzhou). The resulting cell samples were observed under a fluorescence microscope (Olympus, Japan) and imaged with a 10×objective lens. The fluorescence intensity for nucleoli and nucleoplasm in an equal area was obtained with the Image J software. The relative fluorescence intensity for a nucleolus was obtained using the following formula: (the fluorescence intensity of a nucleolus – the fluorescence intensity of nucleoplasm in an equal area to the nucleolus) × the rate of Pol I products in total rRNA (0.983). The scatter plot was achieved by analysing the relative fluorescence intensity of individual nucleolus using the Graphpad Prism 6 software.

### Northern dot blot

HeLa cell lines expressing BRF1 shRNA or HA-BRF1 and their control cell lines were cultured in 10 cm dishes. At 90% confluence, cells were harvested and nuclei were purified from these cells using the method as described above. Next, total RNA was extracted from the nuclei using a RNA extraction kit (Axygen). One μg of total RNA was load in small circles on a piece of Nylon membrane (5 cm × 7 cm), followed by drying at 65℃ for 1 hour. Probes were prepared in a 40 μL reaction mixture containing 10 U klenow enzyme , 25 pmole biotin-labeled random hexamer primer and 500 ng DNA template amplified from the introns of 45S pre-rRNA. Northern hybridization was performed using standard procedures as described previously (*Joseph Sambrook and Russell, 2006*). After hybridization and washing, the membrane was incubated for 1 hour in a 5% skimmed milk-PBS solution containing 1 μL anti-biotin HRP for 1 hour, and it was subjected to washing and detection with ECL reagent.

### In vitro transcription

The rDNA promoter amplified from human genomic DNA was cloned into the pGL3-basic reporter vector. HeLa cells were cultured in 10-cm dishes. At 90% confluence, cells were harvested and nuclei were purified using the method as described above. The resulting nuclei were re-suspended in 400μL buffer D (20 mM HEPES 7.9, 20% Glycerol, 0.1 M KCl, 0.2 mM EDTA, 0.5 mM DTT, 0.5 mM PMSF) and subjected to vortexing for 20 min. Nuclear extract was obtained after centrifuging for 10 min at 12000 rpm. *In vitro* transcription assays were performed using the rDNA promoter-driving plasmids and the resulting nuclear extract according to the nonradioactive method as described previously (*Wang et al., 2015*). The relative transcription activity of the rDNA promoter was obtained by comparing the promoter activity in the nuclear extract with and without BRF depletion.

### Cell proliferation assays

Cell proliferation assays for the cell lines expressing BRF1 shRNA or HA-BRF1 were performed using different methods, including cell counting, MTT and EdU assays. The MTT assay is a colorimetric assay for measuring cell metabolic activity, which is based on the conversion of MTT into formazan crystals by living cells. For the assays with MTT methods, cell proliferation assays were performed as described previously (*Peng et al., 2020*). For the assays with EdU , we followed the manual provided by the BeyoClick EdU Cell Proliferation Kit (Beyotime, China). Briefly, HeLa cells expressing control shRNA or BRF1 shRNA were seeded in 24-well plates. After 24 hours, culture medium was replaced with a complete medium containing 10 mM EdU and cells were incubated for 2 hours. After incubation, cells were fixed for 15 minutes using 4% paraformaldehyde, followed by permeabilizing for 10 minutes with 1.3 % triton-100 PBS solution and staining for 30 minutes using a Click Additive Solution. After which, cell samples were washed 3 times with 1×PBS solution and stained for 10 minutes using the 1×Hoechst-33342 solution. After drying, the samples stained were imaged under a fluorescence microscope (Olympus IX71-F22FL/DIC, Japan). EdU-labeled cells and total cells were counted from minimal five images for each sample. The rate of EdU positive cells was obtained by comparing the number of the EdU-labeled cells to that of the total cells.

### Mouse models for tumour formation

Twenty of five-week-old BALB/C female nude mice were purchased from the Vital River Laboratory Animal Technology Co. (Beijing, China). The mice were housed under Specific Pathogen Free (SPF) conditions in a temperature-controlled room (25 °C) with a 12 h light/dark cycle and 60 % humidity. At six weeks old, the nude mice were randomly grouped into two (10 mice/each group) and were subjected to subcutaneous injection on the right sides of their backs near the hind legs. Each mouse was injected with 1×10^7^ HeLa cells expressing control shRNA or BRF1 shRNA. At 7 days postinjection, the sizes of tumours were measured by Vernier caliper every 3 days; the tumour volume was calculated using the following formula: 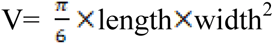. At the end of the fifth week, mice were euthanized. The tumours grown in the mice were removed, weighed and photographed. Immunohistochemistry and hematoxylin and eosin (H&E) staining were performed using the tumour tissues. Animal experiments were approved by the Animal and Medical Ethics Committee of School of Life Science and Health at Wuhan University of Science and Technology. The experiments were conducted according to the Animal Welfare Guidelines (NIH).

### Hematoxylin and eosin staining and immunohistochemistry

Three tumour samples randomly picked from each group were fixed with formalin, embedded with paraffin and sectioned manually according to a standard procedure. Tissue sections were deparaffinized before Hematoxylin and eosin (H &E) staining was performed as described previously (*Canene-Adams, 2013*). The section specimens were observed and imaged under a bright light microscope. For immunohistochemistry (IHC) assays, the paraffin-embedded tissue sections obtained above were deparaffinized twice with xylene and then rehydrated using different concentrations of ethanol and double-distilled water (ddH_2_O). IHC assays were performed using the anti-BRF1 antibody based on the method described previously (*Furlan-Magaril, 2009*). The specimens from IHC were observed and imaged under a bright light microscope with 400x amplification.

### Immunoprecipitation and immunoblotting

HeLa cells were cultured in six 10-cm dishes. At 90% confluence, cells were harvested and nuclei were isolated from the cells using buffer A (10 mM Hepes pH7.9, 10 mM KCl, 1.5 mM MgCl2, 0.5 mM DTT) and a Dounce homogenizer. After centrifugation, nuclei pellet was suspended in 1 mL lysis buffer (50 mM Tris-HCl pH7.9, 150 mM NaCl, 1 mM DTT, 1% NP40%, 2 mM PMSF) and then vortexed for 20 minutes. Nuclear extract was remained for immunoprecipitation (IP) assays after centrifuging 10 minutes at 13000 rpm. IP assays were performed using normal IgG, BRF1 antibody and the antibodies against UBF (ab244287, Abcam), TIF-1A (ab251933, Abcam), TBP (ab28175, abcam), TAF1A (SC-393600, Santa Cruz Biotech), GTF3C2 (SC-81406, Santa Cruz Biotech), TAF1B (CSB-PA684476ESR1HU, CUSABio) and RPA43 (Ab99305, Abcam), respectively. Briefly, 200 μL nuclear extract was incubated with 5 μg antibody at 4℃ overnight, followed by incubating with 40μL protein A/G agarose (Santa Cruz Biotech) at room temperature for 2 h. After incubation, samples were subjected to washing 4 for time using a modified RIPA buffer (0.05%SDS, 0.1% sodium deoxycholate, 1% Triton X-100, 1 mM EDTA, 0.5 mM EGTA, 250 mM NaCl, 10 mM Tris-HCl, Ph8.0). The antibody-bound proteins were eluted into 40 μL 1*SDS loading buffer. Ten μL of the eluates were used for Western blot analysis, where 5% input was used as a positive control.

### Reporter assays and ChIP assays

A cDNA fragment encoding the 5’ prime region (about 400 nt) of 45S rRNA was amplified by RT-PCR and was cloned downstream of the rDNA promoter at the plasmid pGL3-rDNAP (see Fig 7F). HeLa cell lines stably expressing BRF1 shRNA or control shRNA were seeded in 12-well plates and cultured overnight prior to transfection. The rDNA promoter-driving reporter vectors (1.5 μg/well) were transfected into these cell lines with triplicates for control or treatment cell lines. At 48 hours post-transfection, cells were harvested and reporter gene expression was analysed by RT-qPCR using a primer from a luciferase gene (*Figure 7C*) or from 45S pre-rRNA (*Figure 7F*).

ChIP assays were performed using HeLa cell lines stably expressing BRF1 shRNA or HA-BRF1 and their control cell lines as described previously (*Peng et al., 2020*), where BRF1, TBP, TIF-1A, TAF1A, UBF and RPA43 antibodies were used in the assays. DNA was recovered from ChIP samples after chromatin de-crosslinking using a Qiagen PCR purification kit and eluted into 40 μL ddH_2_O. One μL sample from a ChIP assay was used for a qPCR reaction, while 1 ng genomic DNA (equal to 0.03% input) was used as a positive control. Relative occupancy was obtained by comparing the enrichment of promoter DNA in 1 μL ChIP sample to that in 0.03% input. Sequential ChIP assays were performed according to the method as described previously (*Furlan-Magaril et al., 2009*). Briefly, HeLa cells were cultured in 10-cm dishes; at 90% confluence, cells were fixed and harvested for ChIP assays using BRF1 antibody. After eluting from the first immunoprecipitation, the chromatin was diluted and incubated with the antibodies against the Pol I transcription machinery factors. When the second ChIP assays were finished, the DNA was purified as done for the normal ChIP assays. Quantitative PCR was performed using 1 μL sample from the sequential ChIP assays where 0.5 ng genomic DNA (0.01% input DNA) was used as a positive control. All antibodies used as ChIP assays were the same as those for co-IP assays.

### Experimental design and statistical analysis

The experiments, including IF assays, RT-qPCR, cell proliferation assays, CHIP assay and reporter assays were designed and performed in three biological replicates (n=3). The mean, standard deviation (SD) and *p* values for RT-qPCR, cell proliferation assays, tumour growth experiments, reporter assays, and ChIP assays were calculated with the GraphPad Prism 6.0 software. The values in the point or bar graphs represent the mean±SD of three biological replicates. *P* values were obtained by one way ANOVA.

## Acknowledgements

N/A

## Funding information

This work was funded by the project (31671357 to WD, 62172312 to S Zhang) from the Natural Science Foundation of China and by the project from the China Postdoctral Science Foundation (2020M672424 to JW).

## Competing interests

The authors declare no competing interests.

## Abbreviation

BRF1: TFIIB-related factor 1
Pol I: RNA polymerase I
Pol III: RNA polymerase III
rDNA: The DNA encoding 45S ribosomal RNA
FLNA: Filamin A
NPM1: nucleophosmin 1
FBL: Fibrillarin
TBP: TATA-binding protein
TIF-1A: transcription initiator factor 1A
RPA40: DNA-directed RNA polymerase I and III 40 kDa polypeptide
RPA43: DNA-directed RNA polymerase I subunit F
UBF: upstream binding factor
SL1: Selectivity factor 1
DAPI: 4′,6-diamidino-2-phenylindole
RT-qPCR: reverse transcription coupled quantitative polymerase chain reaction
Ctrl IgG: Control immunoglobulin G
GAPDH: glyceraldehyde-3-phosphate dehydrogenas
IF: immunofluorescence
IP: immunoprecipitation
ChIP: chromatin immunoprecipitation
HA: Hemagglutinin
GSK3B: glycogen synthase kinase 3 beta
BDP1: B Double Prime 1
TFIIIB: transcription factor III B
TFIIIC: transcription factor III C
TAF1A: TATA box binding protein associated factor, RNA polymerase I subunit A
TAF1B: TATA box binding protein associated factor, RNA polymerase I subunit B
ICE: internal core element
A (*Figure 8I*): Box A element
C (*Figure 8I*): Box C element
RNA: ribonucleic acid
DNA: deoxyribonucleic acid
Pre-RNA: ribosomal ribonucleic acid precursor
Ub: ubiquitin
EU: 5-ethynyl uridine
EdU: 5-ethynyl-2’-deoxyuridine
CCK-8: Cell counting kit-8.

## Ethics statement

Animal experiments were approved by the Animal and Medical Ethics Committee of School of Life Science and Health at Wuhan University of Science and Technology. The animal protocols abided by the Animal Welfare Guidelines (NIH).

---

**Figure 1-figure supplement.**
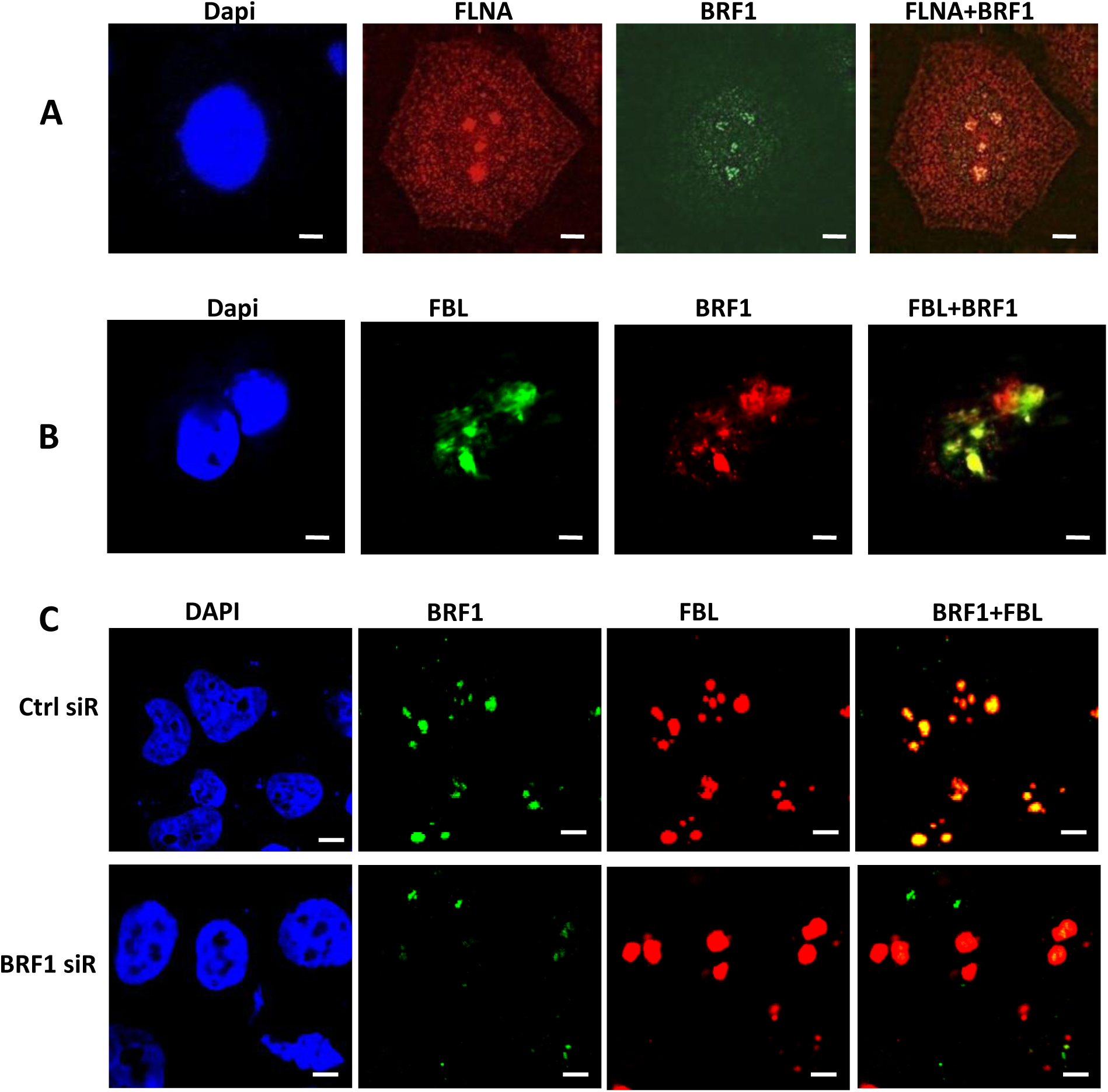
BRF1 is present in the nucleoli within SaOS2 cells. (A)) BRF1 and FLNA were co-localized in SaOs2 cells’ nucleoli. (*B*) BRF1 and fibrillarin were co-localized in SaOS2 cells’ nucleoli. IF assays were performed using SaOS2 cells and the antibodies against BRF1, FLNA, and FBL, respectively. The specimens in A and B were observed under a Decon Vision fluorescence microscopy and imaged with a 60x objective lens (Olympus). The scale bars in A and B represent 1 μm. (*C*) BRF silence reduced BRF1 localization to HeLa nucleoli. IF assays were performed using HeLa cells expressing BRF1 siRNA or control siRNA and the antibodies against BRF1 and FBL, respectively. The specimens were observed under a confocal fluorescence microscopy and imaged with a 60x objective lens (Olympus). The scale bars in C represent 1 μm.

**Figure 2-figure supplement 1.**
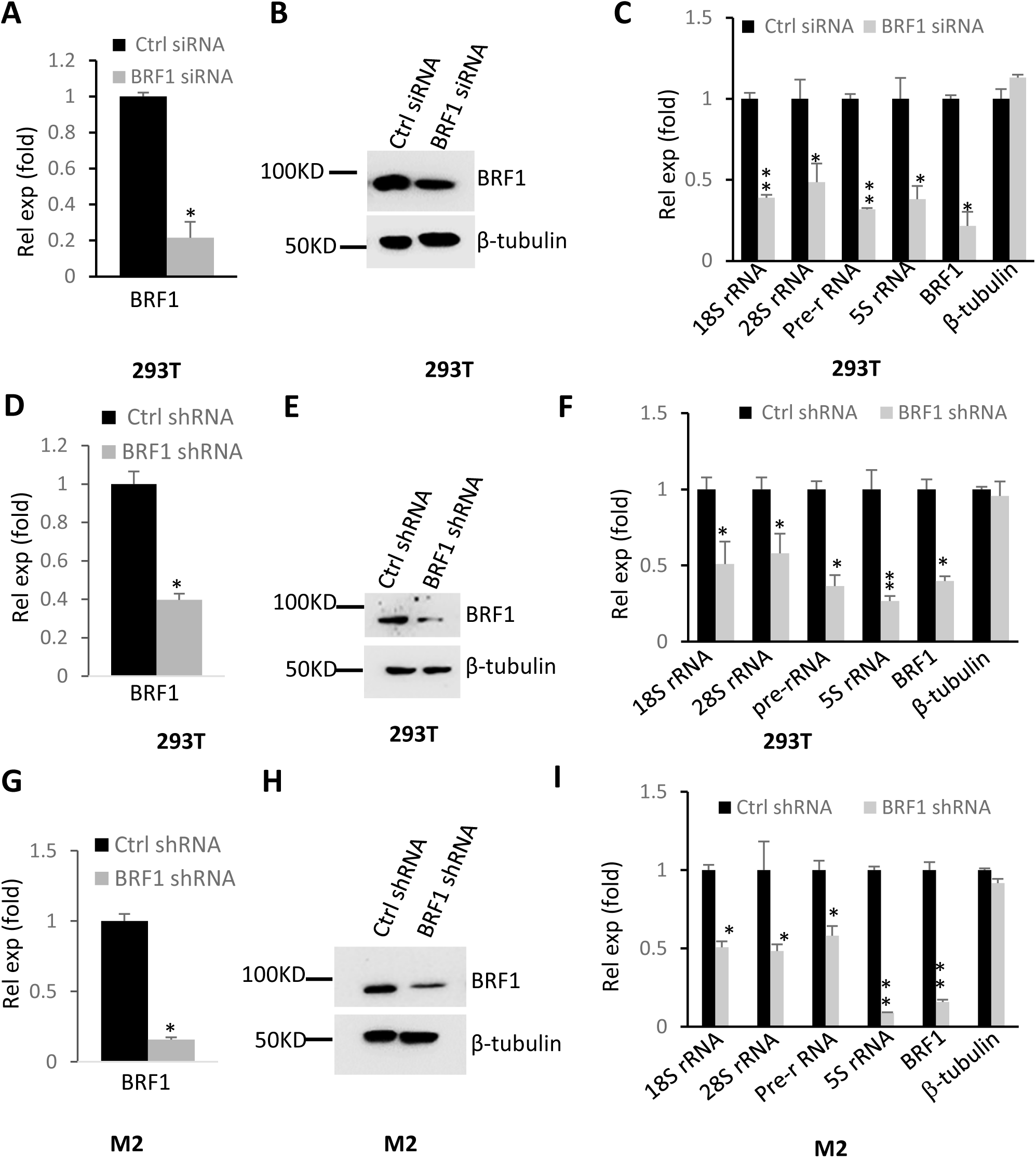
BRF1 knockdown reduced Pol I-mediated transcription. (*A-C*) Transfection of BRF1 siRNA reduced Pol I-directed transcription in 293T cells. 293T cells were transfected with BRF1 siRNA or control siRNA; after 48h, BRF1 mRNA and protein expression levels were detected by RT-qPCR (*A*) and Western blot (*B*), respectively. Ribosomal RNA gene expression was analyzed by RT-qPCR (*C*). (*D-F*) Stable expression of BRF1 shRNA reduced Pol I-directed transcription in 293T cells. 293T cell lines stably expressing BRF1 shRNA or control shRNA were generated using a lentiviral infection system. BRF1 mRNA and protein levels were analyzed by RT-qPCR (*D*) and Western blot (*E*), respectively. Ribosomal RNA gene expression was examined by RT-qPCR (*F*). (*G-I*) BRF1 knockdown reduced Pol I-dependent transcription in M2 cells. M2 cell lines stably expressing BRF1 shRNA or control shRNA were generated using a lentiviral infection system. Expression of BRF1 mRNA (G) and protein (*H*) was detected by RT-qPCR and Western blot, and Ribosomal RNA gene expression (*I*) was analyzed by RT-qPCR. Each column in the bar graphs represents the mean±SD of three biological replicates. *, *P*≦0.05; **, *P*≦0.01, *P* values were obtained by one way ANOVA

**Figure 2-figure supplement 2.**
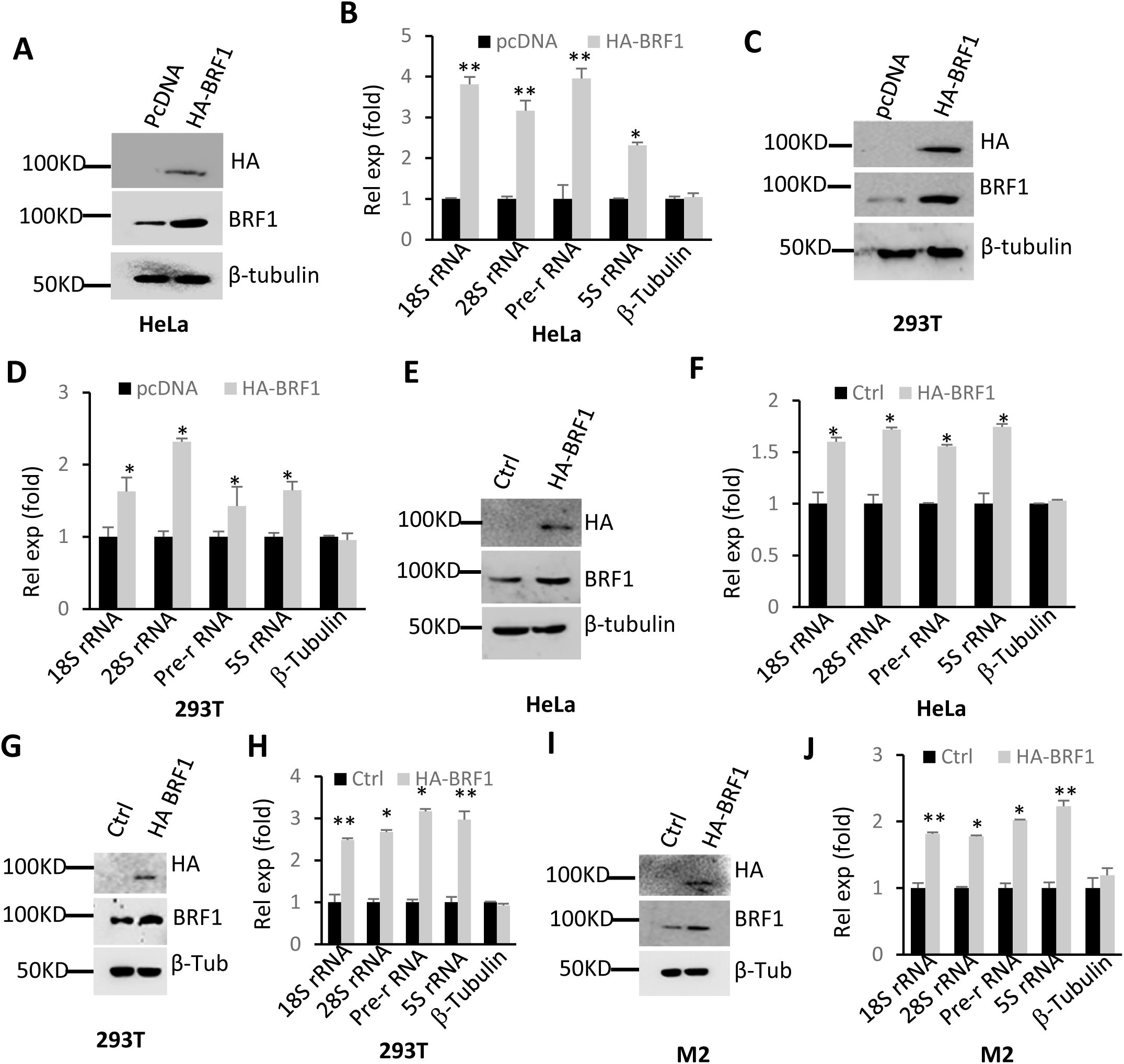
Effect of BRF1 overexpression on Pol I–directed transcription. (*A* and *B*) BRF1 overexpression in HeLa cells stimulated Pol I–directed transcription. HeLa cells were transfected with the expression vectors as indicated. Expression of BRF1 and rRNA gene was monitored by Western blot (*A*) and RT-qPCR (*B*), rescpectively. (*C* and *D*) BRF1 overexpression in 293T cellls enhanced Pol I-dependent transcription. 293T cells were transfected with the expression vectors as indicated. BRF1 protein (*C*) and rRNA levels (*D*) were examined as for *A* and *B*. (*E* and *F*) Stable expression of HA-BRF1 in HeLa cells activated Pol I-directed transcription. A HeLa cell line stably expressing HA-BRF1 and its control cell line were generated and determined by Western blot (*E*). Ribosomal RNA expression was analysed by RT-qPCR (*F*). (*G* and *H*) Stable expression of HA-BRF1 enhanced Pol I-directed transcription in 293T cells. A 293T cell line stably expressing HA-BRF1 and its control cell line were generated as done for HeLa cell lines. HA-BRF1 expression (*G*) and Pol I transcription (*H*) was examined using Western blot and RT-qPCR, respectively. (*I* and *J*) Stable expression of HA-BRF1 increased Pol I-mediated transcription in M2 cells. A M2 cell line stably expressing HA-BRF1 and its control cell line were generated as done for HeLa cells. HA-BRF1 expression (*I*) and Pol I transcription (*J*) was examined using Western blot and RT-qPCR, respectively. Each column in B, D, F, H and J represents the mean±SD of three biological replicates. *, *P*≦0.05; **, *P*≦0.01, *P* values were obtained by one way ANOVA

**Figure 2-figure supplement 3.**
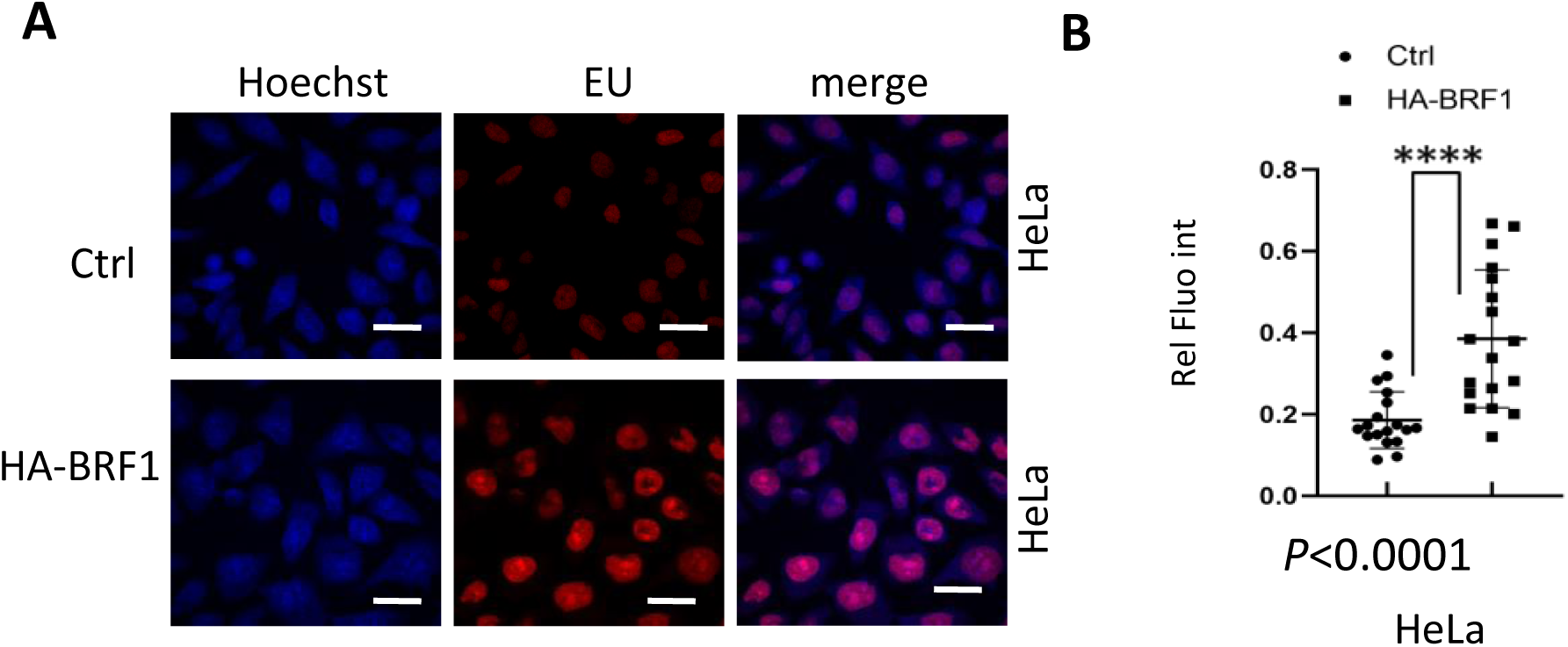
BRF1 overexpression enhanced the rRNA levels newly synthesized in the nulceoli of HeLa cells. A HeLa cell line stably expressing HA-BRF1 and its control cell line were labelled with 5-ethynyl urindine (EU). The EU labelled RNA was detected using a EU detection kit. Images (**A**) were captured under a fluorescence microscope (Olympus) and EU red fluorescence intensity (**B**) within nucleoli and nucleoplasm was obtained using the ImageJ software. The scale bar in each image represents 50 μm.

**Figure 2-figure supplement 4.**
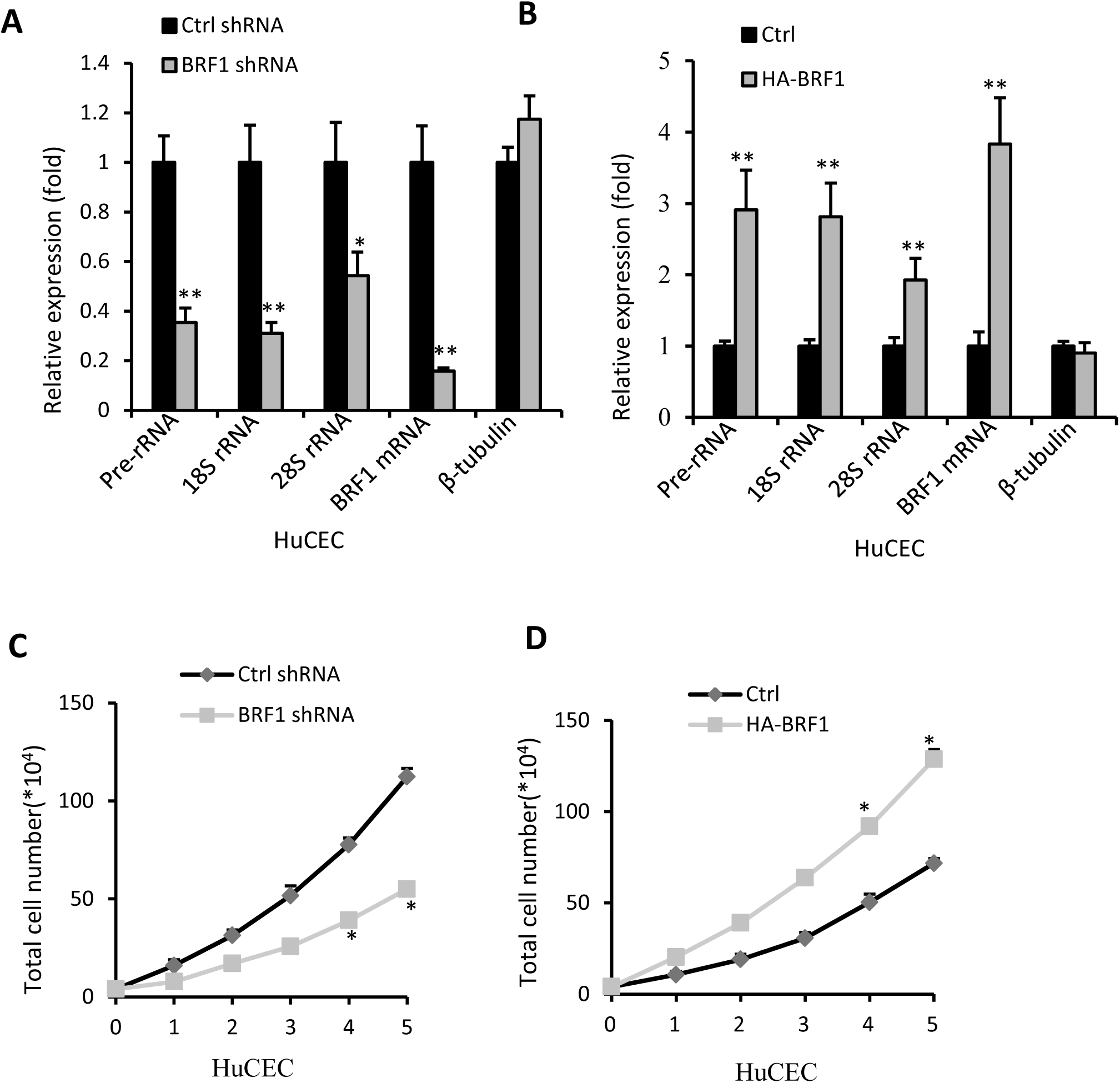
BRF1 promotes Pol I-mediated transcription and proliferation of HuCEC cells. (A) BRF1 silencing decreased expression of Pol I products in HuCEC cells. RT-qPCR was performed using the RNA extracted from HuCEC cells stably expressing BRF1 shRNA or control shRNA. (*B*) BRF1 overexpression increased expression of Pol I products in HuCEC cells. RT-qPCR was performed using the RNA extracted from a HuCEC cells stably expressing HA-BRF1 and its control cell line. (*C*) BRF1 silence reduced HuCEC cell proliferation activity. Cell counting assays were performed using HuCEC cell lines stably expressing BRF1 shRNA or control shRNA. (*D*) BRF1 overexpression increased HuCEC cell proliferation activity. Cell counting assays were performed using a HuCEC cell line stably expressing HA-BRF1 and its control cell line. Each column in the graphs and J represents the mean±SD of three biological replicates. *, *P*≦0.05; **, *P*≦0.01, *P* values were obtained by one way ANOVA

**Figure 4-figure supplement 1.**
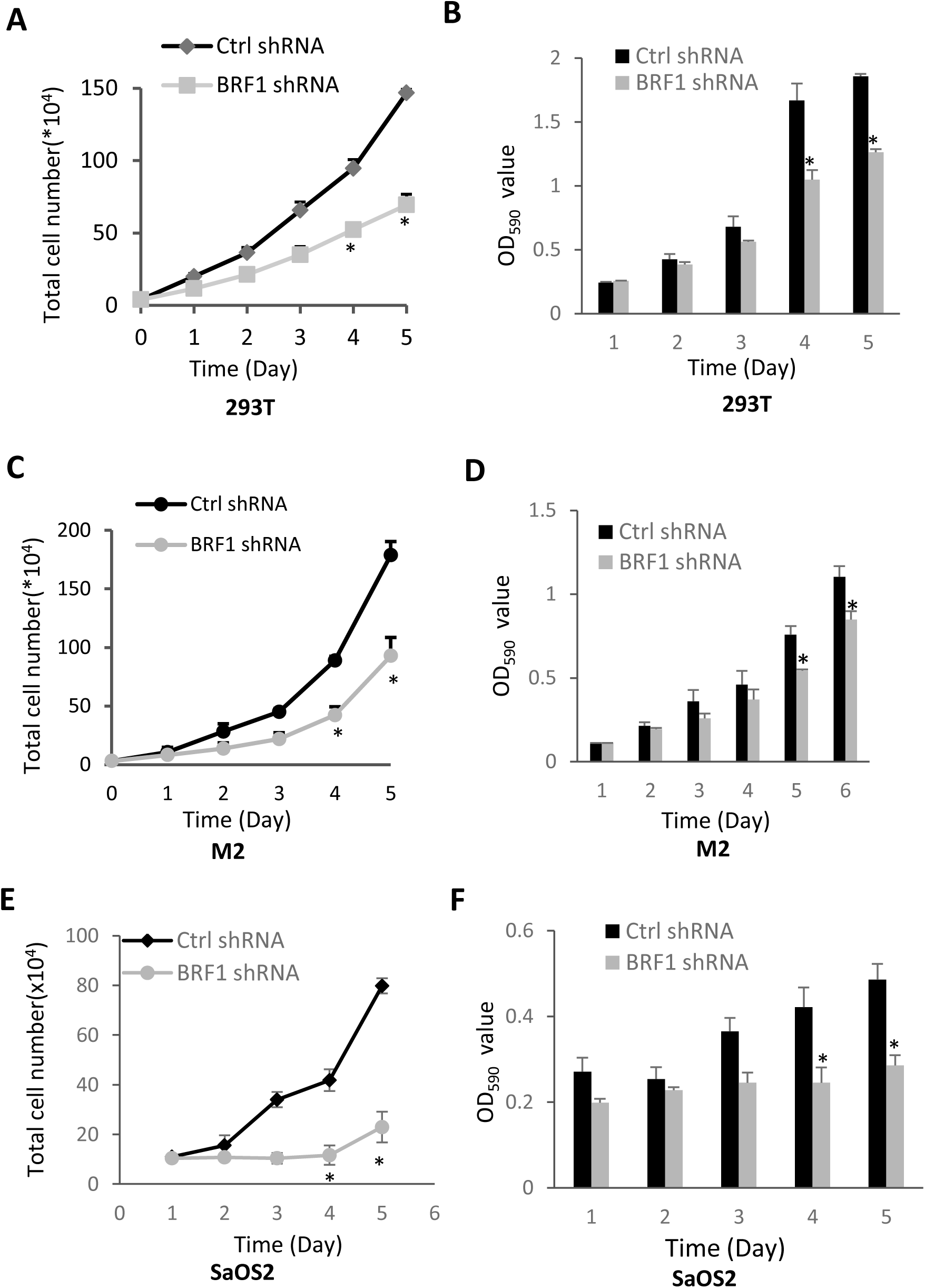
BRF1 knockdown inhibited proliferation activity for transformed cell lines. (*A* and *B*) BRF1 knockdown reduced 293T cell proliferation. (*C* and *D*) BRF1 depletion inhibited M2 cell proliferation. (*E* and *F* ) BRF1 silence repressed SaOS2 cell proliferation. The results in *A*, *C*, and *E* were obtained from cell counting assays, whereas the results in *B*, *D*, and *F* were obtained from MTT assays. Each column in the graphs represents the mean±SD of three biological replicates. *, *P*≦0.05; **, *P*≦0.01. *P* values were obtained by one way ANOVA.

**Figure 4-figure supplement 2.**
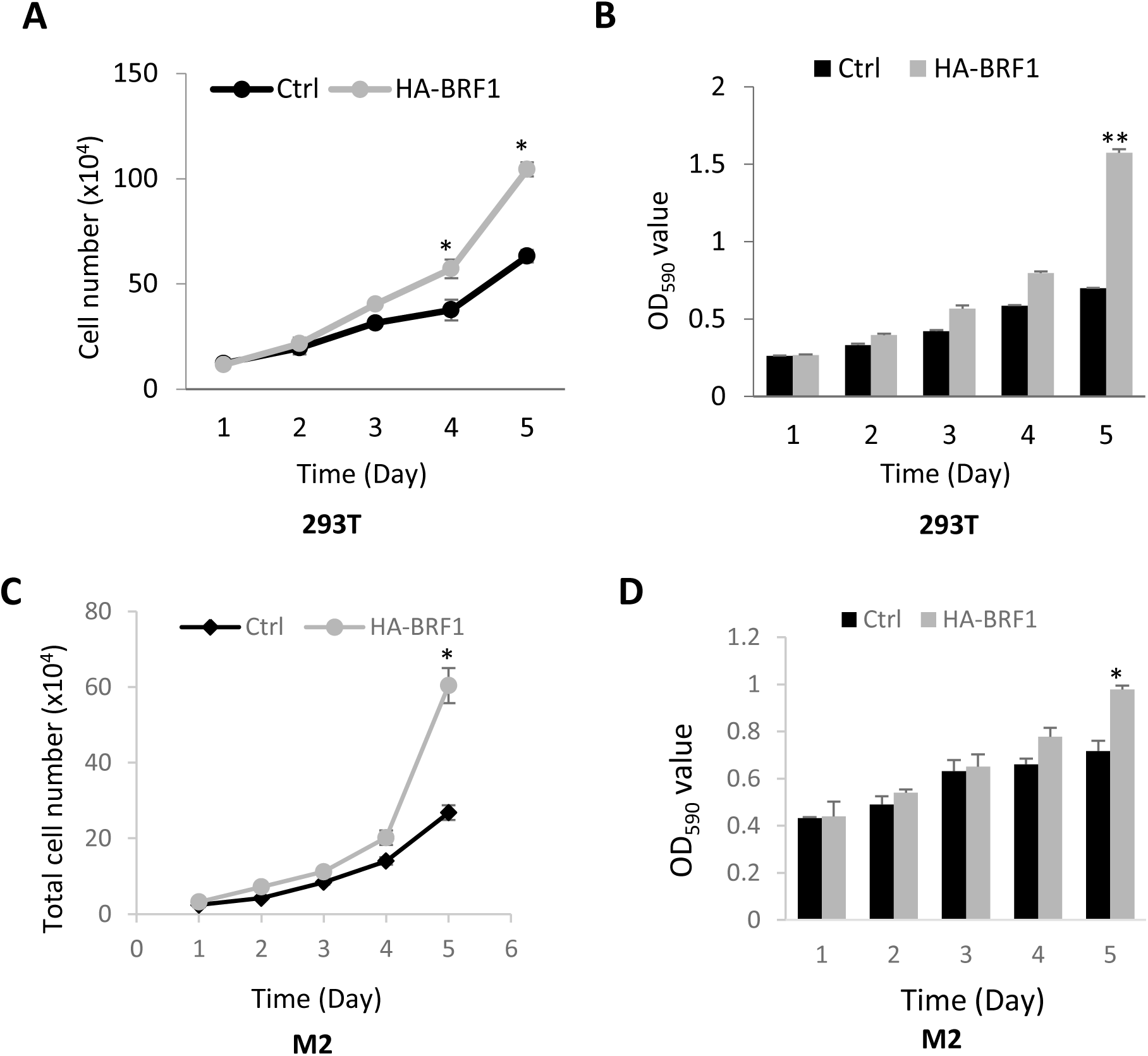
BRF1 overexpression stimulated proliferation activity for transformed cell lines. (*A* and *B*) BRF1 overexpression increased 293T cell proliferation; (*C* and *D*) BRF1 overexpression enhanced M2 cell proliferation. The results in *A* and *C* were obtained from cell counting, assays whereas the results in *B* and *D* were obtained from MTT assays. Each column in the graphs represents the mean±SD of three biological replicates. *, *P*≦0.05; **, *P*≦0.01, *P* values were obtained by one way ANOVA.

**Figure 4 –figure supplement 3.**
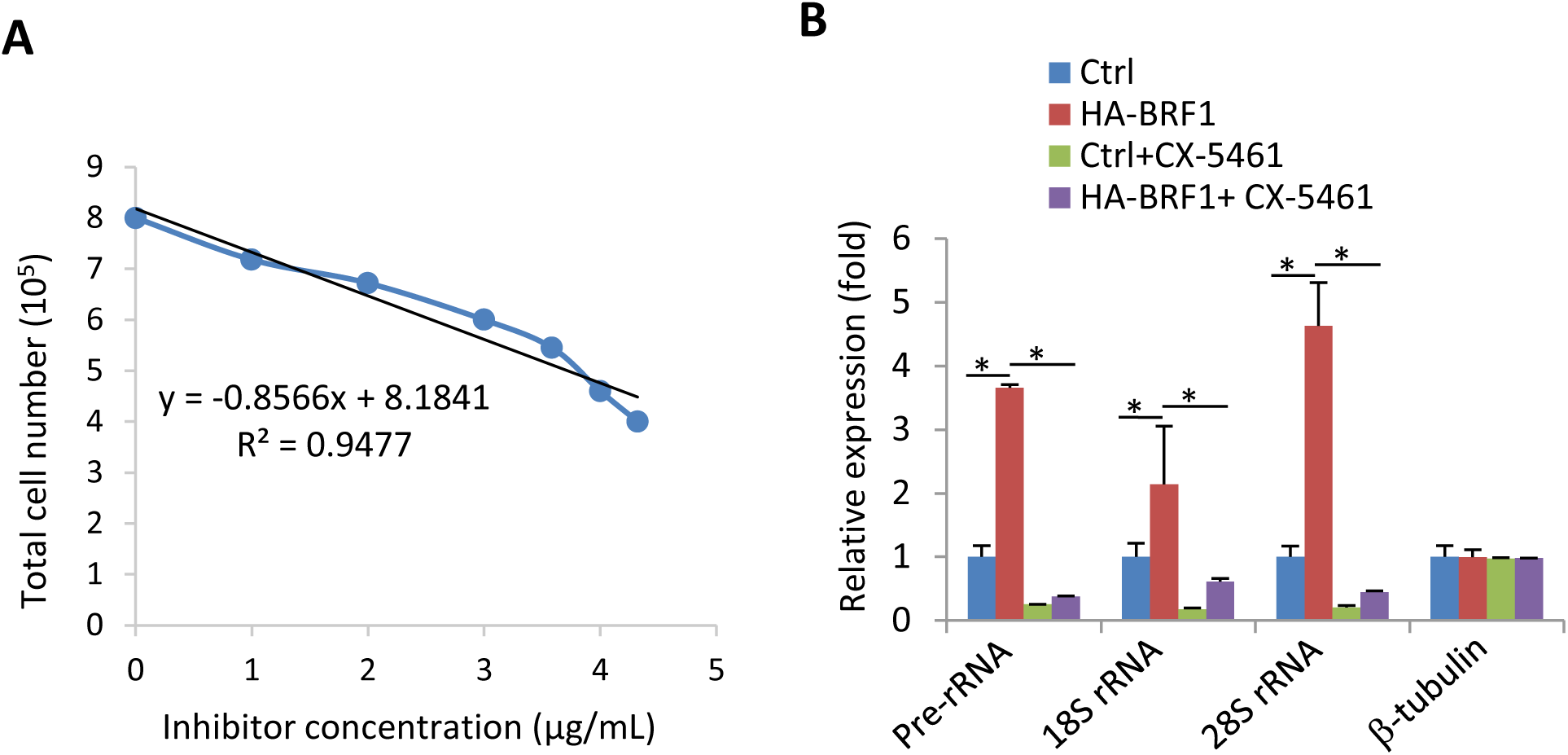
CX-5461 inhibited the activation of Pol I transcription caused by BRF1 overexpression in HeLa cells. (*A*) The analysis of correlation between the concentrations of the inhibitor CX-5461 and the number of HeLa cells. IC50 was determined by the linear regression equation. (*B*) the inhibitor CX-5461 significantly impeded Pol I transcriptional activation caused by BRF1 overexpression. A HeLa cell line stably expressing HA-BRF1 or its control cell line were seeded in 6-well plates and cultured for 24 hours. An inhibitor (CX-5461) was added to the HA-BRF1-expressing cells at the final concentration of 4 μg/mL. The cells were harvested for RT-qPCR assays after incubating for 4 hours in the medium containing the inhibitor. *, *P*≦0.05; **, *P*≦0.01, *P* values were obtained by one way ANOVA.

**Figure 6-figure supplement 1.**
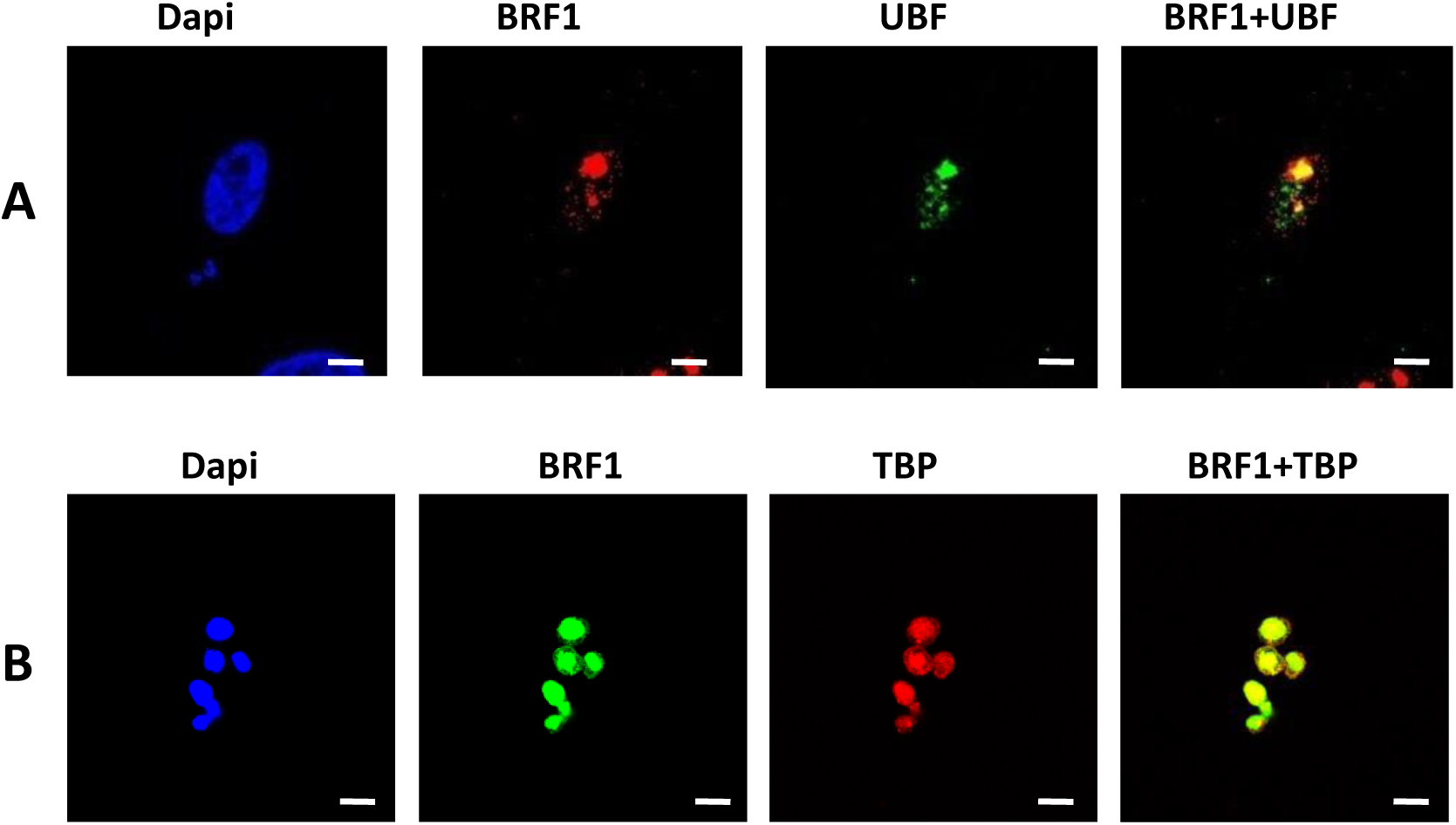
The co-localization analysis between BRF1 and Pol III transcription factors in HeLa cells. (A) BRF1 and UBF were co-localized in the nucleoli of HeLa cells. Immunofluorescence assays were performed using the HeLa cells and the antibodies against BRF1 and UBF, respectively. (*B*) BRF1 and TBP were co-localized in the nucleoli purified from HeLa cells. IF assays were performed using the nucleoli particles purified from HeLa cells and the antibodies against BRF1 and TBP. The scale bars in *A* and *B* represent 5 μm.

**Figure 6-figure supplement 2.**
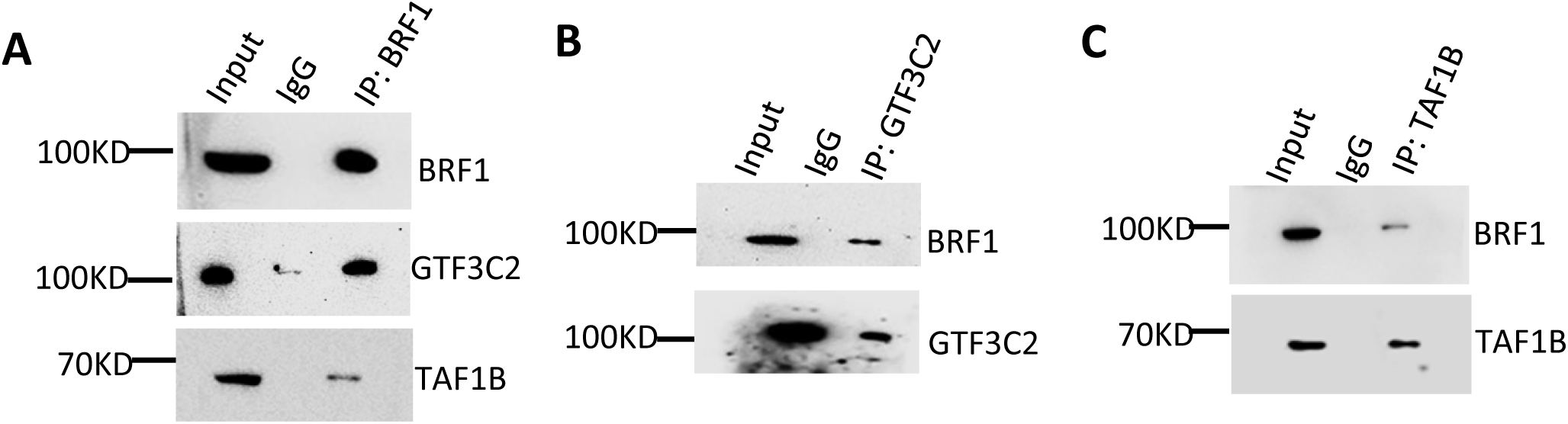
BRF1 binds to TFIIIC subunit GTF3C2 and SL1 subunit TAF1B. (A) BRF1 IP results showing BRF1 association with GTF3C2 and TAF1B. IP assays were performed using BRF1 antibody and HeLa nuclear extract. BRF1-bound protein was detected by Western blot using the antibodies as indicated. (*B*) GTF3C2 IP result showing GT3C2 association with BRF1. IP assays were performed using GTF3C2 antibody and HeLa nuclear extract. GTF3C2-bound protein was detected by Western blot using the antibodies as indicated. (*C*) TAF1B IP result showing TAF1B association with BRF1. IP assays were performed using TAF1B antibody and HeLa nuclear extract. TAF1B-bound protein was detected by Western blot using the antibodies as indicated.

## Notes

### Competing Interest Statement

The authors have declared no competing interest.

